# RNA-dependent chromatin organization during development

**DOI:** 10.64898/2026.08.14.744931

**Authors:** Alexa Alipour, Nidhi Rani Lokesh, Mark E. Pownall

## Abstract

Embryonic development is characterized by controlled spatiotemporal remodeling of the nuclear landscape. To understand how this process is regulated, it is crucial to identify the factors controlling nano and mesoscale nuclear organization. Here, we developed an improved Chromatin Expansion Microscopy approach to visualize nuclear organization in developing zebrafish embryos at nanometer-scale resolution. We observe a stepwise emergence of chromatin compaction during early development, coincident with the onset of heterochromatin formation. Perturbation of the facultative heterochromatin mark H3K27me3 reveals that it is neither necessary nor sufficient for chromatin compaction in vivo. Instead, inhibiting transcription elongation disrupts chromatin compaction, while increasing nuclear RNA levels enhances compaction independently of transcriptional activity. Together, these results identify nuclear RNA as a major determinant of chromatin organization during development and support a model in which RNA promotes mesoscale chromatin compaction independently of canonical heterochromatin pathways.

**One-Sentence Summary:** In vivo nanoscale imaging reveals RNA-dependent chromatin compaction during development.

---

Early embryogenesis is characterized by the rapid establishment of new patterns of chromatin organization that enable the first major cell fate transitions after fertilization (*1–6*). This transition involves the emergence of higher-order nuclear architecture, including physical chromatin compaction, from an initial state that lacks features of mature genome organization (*1*, *7–13*). Although genome-wide studies have defined the timing of transcriptional and epigenetic remodeling during early development (*3*, *5*, *6*, *14–17*), how these molecular changes translate into the physical organization of chromatin in vivo remains poorly understood, in part due to the difficulty of visualizing nuclear structures at nanoscale resolution in intact embryos (*18*, *19*).

A central question in early development is what drives the emergence of higher-order chromatin organization, including the formation of compacted domains and transcriptionally distinct nuclear compartments. In particular, it is unclear how physical chromatin organization relates to the establishment of transcriptional repression in vivo. The deposition of histone H3 lysine 27 trimethylation (H3K27me3), a hallmark of facultative heterochromatin, is closely associated with both gene silencing and large-scale chromatin reorganization (*20–26*). Although H3K27me3 is associated with compacted chromatin (*22*, *27*, *28*), whether compaction is necessary for silencing remains unclear (*29–31*). Further, how these relationships are established during development, and whether H3K27me3 is required for the emergence of chromatin compaction in vivo, are poorly understood.

Here, we used zebrafish embryos to address these questions because early developmental stages are characterized by a chromatin state that is largely devoid of repressive histone modifications, exhibits minimal compaction, and lacks mature three-dimensional genome organization (*4*, *10*, *11*, *13*, *17*, *32*). To directly visualize the emergence of nuclear organization, we developed an improved Chromatin Expansion Microscopy (ChromExM) (*33*) workflow that enables ∼6,000-fold volumetric expansion of developing zebrafish embryos. Using this strategy, we show that chromatin compaction emerges independently of H3K27me3, which is neither necessary nor sufficient to drive compaction, and instead depends on nuclear RNA as a key determinant of chromatin organization during early development.

## Optimized nanoscale imaging reveals stepwise chromatin compaction during early development

ChromExM has provided a powerful tool to visualize nanoscale nuclear organization; however, the first generation technology was low-throughput and required semi-manual analysis, effectively restricting its use to single developmental stages (*33*). To overcome these challenges, we developed an improved ChromExM workflow to enable quantitative analysis of chromatin organization across development by streamlining sample preparation and developing machine learning approaches for downstream analyses. This improved method also increased the linear expansion factor from ∼15-fold to ∼18-fold (mean linear expansion factor, 18.6x) (Fig. S1A to G; see Materials and Methods). Together, these improvements enable direct measurement of chromatin organization across developmental stages within intact embryos (Fig. 1A), allowing high-resolution stage-resolved single-cell analysis of chromatin in vivo.

**Figure 1.**
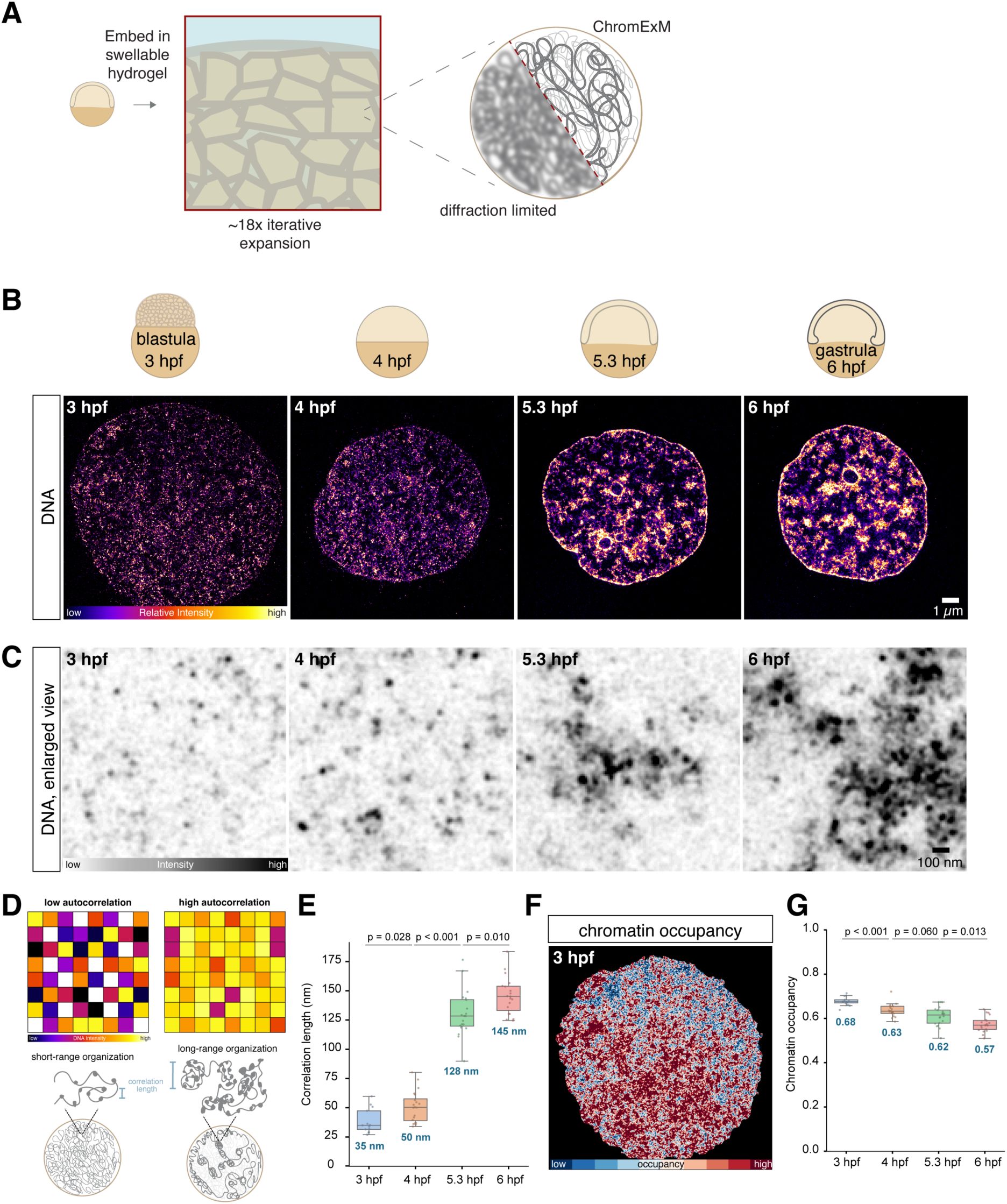
Improved ChromExM reveals emergent nanoscale chromatin organization. (**A**) Schematic of the ChromExM workflow used in the study. (**B**) ChromExM images of individual nuclei from 3, 4, 5.3, and 6 hours post-fertilization (hpf) embryos showing the gradual emergence of chromatin organization. Images shown as single Z-slices; display settings are not matched across images. N = 13, 20, 20, and 20 nuclei from 2 embryos at each stage, respectively. (**C**) Zoomed in views from the images shown in (B) showing nucleosome-scale organization at 3 and 4 hpf, and mesoscale compaction at 5.3 and 6 hpf. (**D**) Schematic of autocorrelation length measurement showing how correlation length may relate to local chromatin organization. (**E**) Quantification of the autocorrelation length from (B) shows a significant increase in correlation length across each developmental stage. P-values determined by permutation test. (**F**) Visualization of chromatin occupancy measurement at 3 hpf. Red corresponds to higher chromatin occupancy and blue corresponds to lower chromatin occupancy. A single Z-slice is shown. (**G**) Quantification of chromatin occupancy as a proportion of nuclear volume shows decreasing chromatin occupancy as chromatin compacts from 3-6 hpf. P-values determined by permutation test.

Using this new strategy, we imaged the DNA in zebrafish embryos at 3, 4, 5.3, and 6 hours post fertilization (hpf), spanning the developmental transition from zygotic genome activation (ZGA) and totipotency to the onset of gastrulation as cells begin to differentiate. This revealed a stepwise emergence of chromatin compaction as embryos undergo early differentiation (Fig. 1B and C; Fig. S1G). At 3 and 4 hpf, we observed DNA signal throughout the nucleus with many small puncta of intense signal (Movie S1). In contrast, at 5.3 and 6 hpf, we observed a high degree of compartmentalization in the nucleus; there are large clusters of intense DNA staining among pockets of low or no signal (Fig. 1B and C; Movie S2). To quantify these changes, we analyzed the radial autocorrelation of the DNA signal, which we used to measure the length scales over which DNA intensity is correlated. We expect that short-range DNA organization will be reflected by short correlation lengths, while long-range organization will be associated with longer correlation lengths (Fig. 1D). At 3 and 4 hpf, chromatin exhibits relatively short autocorrelation lengths (∼35-65 nm), which are on the scale of nucleosomes or nucleosome clutches (*34*) (Fig. 1E; Fig. S1H). This is consistent with the absence of higher-order chromatin structure during ZGA (*10*, *13*, *32*) and the bright puncta observed in the images (Fig. 1C). In contrast, at 5.3 and 6 hpf, DNA displays a significantly longer autocorrelation length (∼130-150 nm) (Fig. 1E; Fig. S1H). We expect this reflects long-range chromatin organization which emerges at the same developmental stages (*13*, *32*). These results reveal a progressive transition that begins with relatively homogeneous chromatin organization containing only short-range structure, consistent with local nucleosome-scale organization. Over time, chromatin becomes spatially heterogeneous, adopting a domain-like architecture, showing the emergence of mesoscale chromatin organization during early development.

As an independent measure of chromatin organization, we next quantified chromatin occupancy, which we defined as the proportion of the nuclear volume containing DNA (Fig. 1F). We observed that chromatin occupancy significantly decreases over development, from ∼68% at 3 hpf to ∼57% at 6 hpf (Fig. 1G). This result is consistent with the formation of compacted chromatin domains and increased interchromatin space. Together with the increasing autocorrelation length, these results demonstrate that mesoscale chromatin organization emerges progressively during early development.

## H3K27me3 is not required for mesoscale chromatin compaction

We next sought to identify the molecular mechanisms that drive this emergent chromatin compartmentalization. The establishment of facultative heterochromatin marked by H3K27me3 is closely associated with chromatin compaction and gene silencing during development, making it a strong candidate regulator of this process (*20–26*). To test whether H3K27me3 contributes to this transition, we first examined its spatial and temporal relationship with chromatin organization. Using diffraction-limited immunostaining, we observed that H3K27me3 levels increase progressively from 4 to 6 hpf, coinciding with the emergence of chromatin compaction (Fig. 2A; Fig. S2A to C). Using one round of expansion (4.5x linear expansion factor; Fig. S2D; see Materials and Methods), we visualized H3K27me3 and DNA at higher spatial resolution and observed that many compacted chromatin sites were enriched with H3K27me3 (Fig. 2B). These observations confirm that H3K27me3 is associated with compact chromatin domains, but do not establish whether it plays a causal role in their formation.

**Figure 2.**
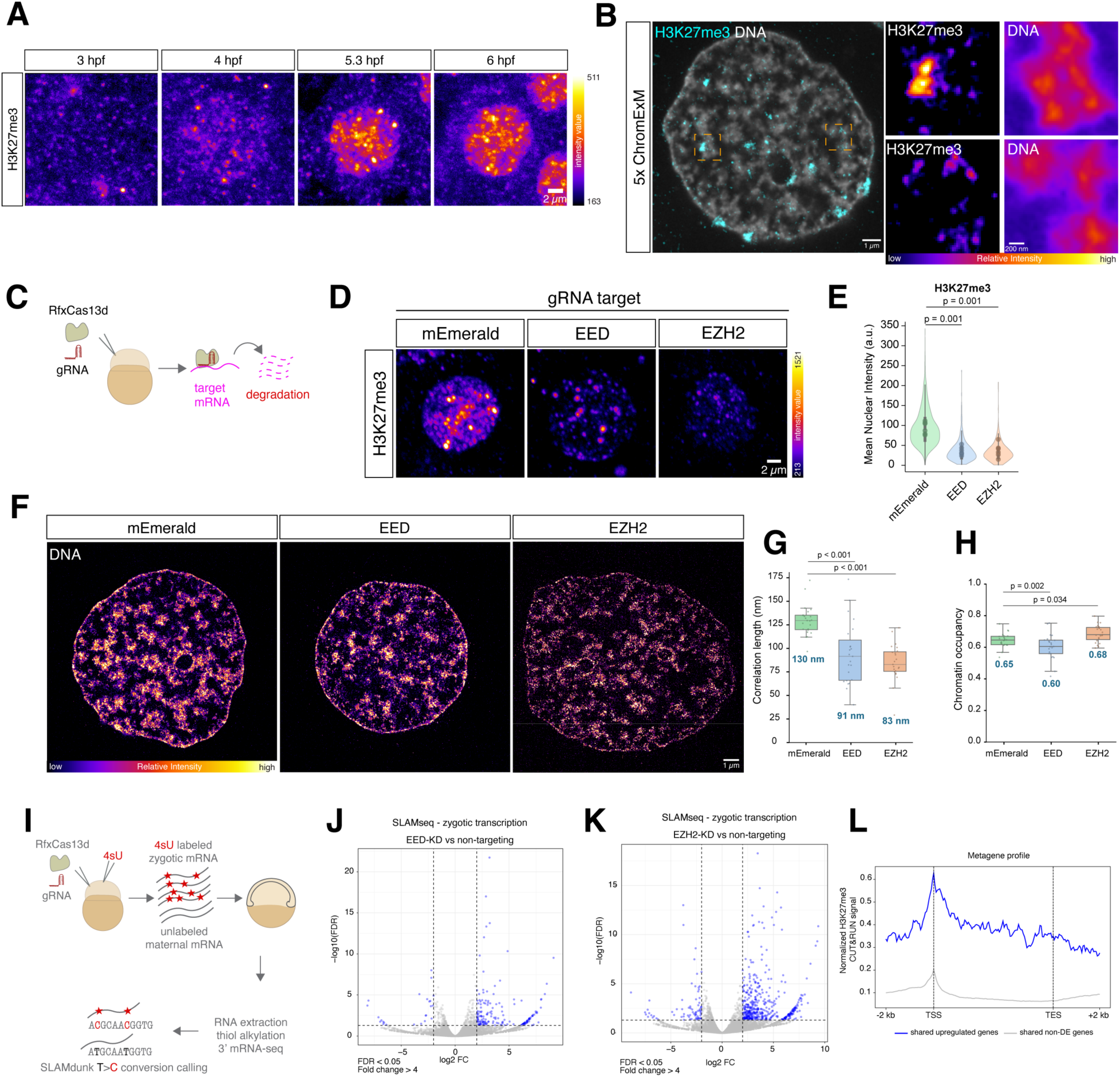
H3K27me3 is dispensable for mesoscale chromatin compaction but required for developmental gene silencing. (**A**) Diffraction-limited immunostaining of H3K27me3 shows gradual accumulation from 3-6 hpf. Images are maximum intensity projections of individual nuclei. N = 103, 291, 825, and 1,077 nuclei at 3, 4, 5.3, and 6 hpf, respectively, from 4 embryos at each stage. (**B**) Once expanded ChromExM of H3K27me3 and DNA at 6 hpf. Insets are of the boxed regions. Images shown as single Z-slices. N = 10 nuclei from 2 embryos. (**C**) Schematic of Cas13d knockdown strategy targeting core components of PRC2, EZH2 and EED. (**D**) Diffraction-limited immunostaining of H3K27me3 in mEmerald-, EED-, and EZH2-targeted embryos at 6 hpf. Images are maximum intensity projections of individual nuclei. N = 1,326, 576, and 424 nuclei for mEmerald-, EED-, and EZH2-targeting, respectively, from 6 embryos in each condition. (**E**) Quantification of nuclear H3K27me3 levels from (D) shows a significant decrease upon PRC2 knockdown. P-values determined by permutation test. (**F**) ChromExM images of nuclei following Cas13d targeting of EZH2 or EED when compared to a non-targeting control (mEmerald). N = 23 nuclei from 2 embryos in each condition. (**G**) Quantification of the autocorrelation length from (F) shows a significant decrease in correlation length upon PRC2 perturbation. P-values determined by permutation test. (**H**) Quantification of chromatin occupancy shows minimal chromatin decompaction upon PRC2 knockdown. P-values determined by permutation test. (**I**) Schematic of SLAM-seq experiment for maternal and zygotic RNA sequencing. (**J**) Volcano plot showing differentially expressed genes in EED-knockdown embryos. (**K**) Volcano plot showing differentially expressed genes in EZH2-knockdown embryos. (**L**) Metaplot of H3K27me3 CUT&RUN signal from 6 hpf embryos (*43*) showing that genes upregulated following EZH2 and EED knockdown are enriched for H3K27me3 relative to unaffected genes.

To directly test whether H3K27me3 is required for chromatin compaction, we disrupted the activity of its writer complex, Polycomb Repressive Complex 2 (PRC2), using CRISPR-RfxCas13d (Cas13d) to degrade the mRNA of core PRC2 components (*21*, *35–37*) (Fig. 2C). We targeted EZH2 or EED mRNAs, two core components of PRC2, for acute degradation and used non-targeting gRNAs against mEmerald or mCherry transcripts, which are not expressed in embryos, as controls. This resulted in an 87% and 81% reduction in EED and EZH2 transcript levels, respectively (Fig. S2E and F). Immunostaining in PRC2-knockdown embryos showed a ∼60-61% reduction in H3K27me3 levels (Fig. 2D and E; Fig. S2G).

To assess how this reduction in H3K27me3 affects chromatin organization, we next performed ChromExM on PRC2-targeted embryos (Fig. 2F; Fig. S2H). Quantification revealed that the autocorrelation length was reduced from ∼135 nm to ∼80-90 nm upon PRC2 depletion (Fig. 2G; Fig. S2I). Despite this shift in the length scale of chromatin organization, chromatin occupancy was largely similar across conditions and, in the case of EED-targeting, mildly decreased (mean chromatin occupancy: 0.65, 0.60, 0.68 for mEmerald-, EED-, and EZH2-targeted embryos, respectively) (Fig. 2H). To exclude the possibility that compaction is maintained by a compensatory increase in H3K9me3, another major heterochromatin mark, we performed immunostaining of H3K9me3 in PRC2-knockdown embryos and observed that global levels of this mark were unchanged in response to reduced H3K27me3 (Fig. S2J and K). This suggests that the remaining compaction is not due to compensation by H3K9me3 and indicate that substantial reduction of H3K27me3 does not prevent mesoscale chromatin compaction.

To confirm these results, we independently perturbed H3K27me3 deposition using the EZH2 catalytic inhibitor tazemetostat (*38*, *39*). Immunostaining in EZH2-inhibited embryos showed an 85% reduction in H3K27me3 levels, with no changes in H3K9me3 levels (Fig. S3A to D). ChromExM analysis revealed no significant changes in chromatin autocorrelation length (DMSO = 145 nm vs tazemetostat = 138 nm; P = 0.532) or chromatin occupancy (DMSO = 0.61 vs tazemetostat = 0.64; P = 0.210) relative to control embryos (Fig. S3E to H), indicating that reducing H3K27me3 levels via EZH2 inhibition does not disrupt mesoscale chromatin organization.

Together, these results indicate that H3K27me3 is not a primary determinant of mesoscale chromatin compaction during early development.

## Global chromatin compaction is not sufficient for developmental gene silencing

Given that depletion of H3K27me3 does not impair global chromatin compaction, we next asked whether transcriptional repression depends on H3K27me3 despite the persistence of chromatin compaction. To this end, we performed thiol(SH)-linked alkylation for the metabolic sequencing of RNA (SLAM-seq) (*40–42*) to measure transcriptome-wide changes in zygotic gene expression (Fig. 2I; Fig. S4A-E). We first analyzed both maternal and zygotic transcripts from SLAM-seq to further assess the specificity of our Cas13d targeting approach. Comparing EED- to EZH2-knockdown embryos identified only 38 differentially expressed (DE) genes, with the targeted transcripts, EZH2 and EED themselves, representing the two most significant hits (Fig. S4F). This demonstrates that Cas13d effectively and specifically depletes the targeted transcripts with minimal off-targets. Similarly, comparing zygotic transcript levels from mEmerald-targeted embryos with those from uninjected embryos showed a strong correlation (Pearson r = 0.969) (Fig. S4A), confirming that Cas13d injection alone does not substantially alter the zygotic transcriptome (*37*).

Comparing zygotic transcripts in PRC2-knockdown embryos to control-targeted embryos revealed 200 and 544 DE genes for EED- and EZH2- targeting, respectively (Fig. 2J and K). Among those, the vast majority of affected genes were upregulated following PRC2 depletion (86.8% and 85.5% and for EED- and EZH2- targeting, respectively), with a substantial overlap between perturbations (Fig. 2J and K). In total, 132 upregulated genes were shared between EZH2- and EED-knockdown, representing 75% of genes upregulated following EED depletion. To confirm that upregulated genes are PRC2 targets, we analyzed published H3K27me3 CUT&RUN data (*43*). Metagene analysis revealed strong enrichment of H3K27me3 centered on the transcription start site (TSS) and extending across the gene body for the upregulated genes (Fig. 2L; Fig. SG and H). Quantification of H3K27me3 signal within TSS-proximal regions demonstrated significantly greater occupancy at shared upregulated genes compared to shared non-differentially expressed genes (2.18-fold higher median signal; Mann-Whitney U test, P = 3.6×10⁻¹²).

These changes are consistent with the canonical role of PRC2 and H3K27me3 in transcriptional repression. However, because zygotic transcription was derepressed without loss of chromatin compaction, these results indicate that PRC2-mediated repression and mesoscale chromatin compaction are separable during early development.

## Transcription promotes chromatin compaction

Given that chromatin compaction persists independently of H3K27me3 and is not sufficient to repress transcription, we next asked whether transcription contributes to the establishment of chromatin organization. To test this, we inhibited transcription elongation using α-amanitin, which prevents DNA translocation by Pol II, leading to broad inhibition of Pol II transcription (Fig. S5A and B) (*33*, *44*, *45*). ChromExM revealed a marked loss of chromatin compaction at 6 hpf, with the autocorrelation length reduced by 52% from ∼145 nm to ∼70 nm while chromatin occupancy increased by 28% (Fig. 3A to D; Fig. S5C and D; Movie S3). These changes indicate a loss of mesoscale chromatin organization, with chromatin adopting a more homogeneous, decompacted state in absence of active transcription.

**Figure 3.**
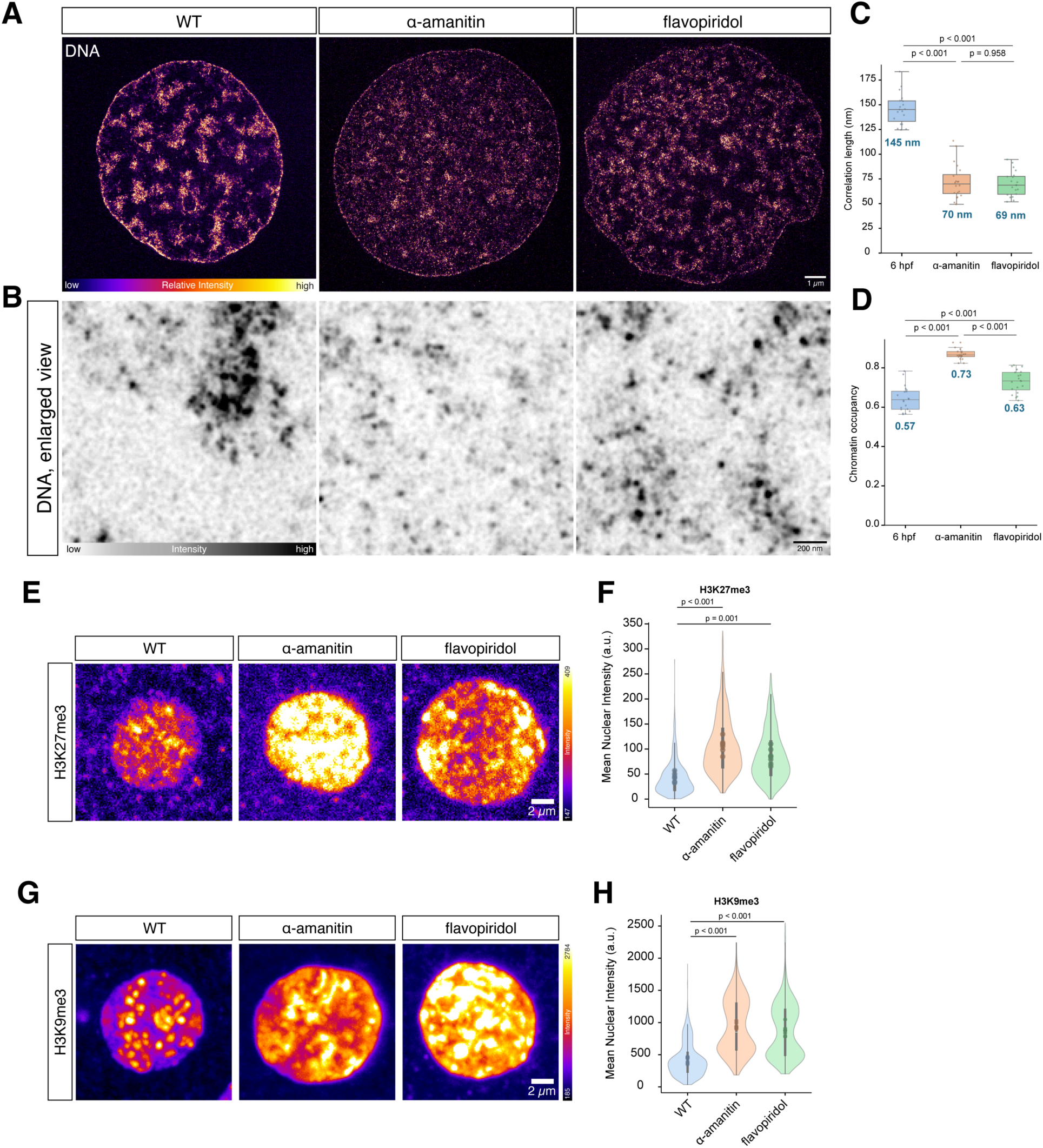
Transcription is required for chromatin compartmentalization. (**A**) ChromExM images of individual nuclei from untreated (wildtype), α-amanitin-, and flavopiridol-treated embryos at 6 hpf showing reduced chromatin compaction when transcription is inhibited. N = 20 nuclei from 2 embryos for each condition. Wildtype images are from the same experiment as those used in Fig. 1B. Images shown as single Z-slices; display settings are not matched across images. (**B**) Zoomed in views from the images shown in (A). (**C**) Quantification of the autocorrelation length from (A) shows significant decrease in correlation upon α-amanitin or flavopiridol treatment. P-values determined by permutation test. (**D**) Quantification of chromatin occupancy from (A) shows a significant increase in chromatin occupancy corresponding to chromatin decompaction upon α-amanitin or flavopiridol treatment. P-values determined by permutation test. (**E**) Diffraction-limited immunostaining of H3K27me3 shows a significant increase following transcription inhibition. N = 2,091, 1,938, and 911 nuclei from 6-7 embryos each for WT, α-amanitin, and flavopiridol, respectively. (**F**) Quantification of nuclear H3K27me3 levels from (E) shows a significant increase upon transcription inhibition. P-values determined by permutation test. (**G**) Diffraction-limited immunostaining of H3K9me3 shows a significant increase following transcription inhibition. N = 2,061, 1,058, and 1,038 nuclei from 8 embryos each for WT, α-amanitin, and flavopiridol, respectively. (**H**) Quantification of nuclear H3K9me3 levels from (G) shows a significant increase upon transcription inhibition. P-values determined by permutation test.

To further dissect the role of transcription in chromatin organization, we next inhibited the Pol II pause release factor CDK9 using flavopiridol to block productive elongation while allowing upstream promoter-proximal transcription to persist (Fig. S5A and B) (*46*, *47*). Under these conditions, chromatin organization was partially disrupted, with reduced autocorrelation length (52%) and a modest increase in chromatin occupancy (∼10%) (Fig. 3A-D; Fig. S5C and D; Movie S4). Compared to α-amanitin treatment, chromatin was more compacted, suggesting that residual transcriptional activity is sufficient to partially maintain chromatin organization (*46*). The distinct effects of α-amanitin and flavopiridol, which inhibit transcription through distinct mechanisms and to different extents, suggest that loss of chromatin organization scales with the degree of transcriptional inhibition and support a specific relationship between ongoing transcription and chromatin organization. Together, these results indicate that ongoing transcription contributes to the establishment of mesoscale chromatin compaction.

Notably, transcription inhibition resulted in a 2-2.5-fold increase in nuclear H3K27me3 levels (Fig. 3E and F; Fig. S5E and F), indicating that chromatin decompaction is not due to a loss of the histone modification. This is consistent with previous observations that transcription inhibition can promote ectopic H3K27me3 accumulation across the genome (*48*). Further, we observed that H3K9me3 levels also increased 2-2.2-fold following transcription inhibition (Fig. 3G and H; Fig S5G). Thus, despite pronounced chromatin decompaction, embryos retain robust accumulation of major repressive histone modifications, indicating that these chromatin regulatory pathways remain active following transcription inhibition. However, the accumulation of canonical heterochromatin marks, even at elevated levels, is not sufficient to establish chromatin compaction in the absence of transcriptional activity. Together, these results indicate that transcription promotes chromatin compaction independently of repressive histone modifications.

## RNA organizes chromatin during development

Because transcription inhibition alters many aspects of cellular physiology downstream of RNA production, we next asked whether a complementary gain-of-function perturbation to directly increase nuclear RNA abundance would be sufficient to alter chromatin organization. To do so, we blocked RNA decay to increase nuclear RNA levels independently of transcription by depleting components of the nuclear RNA exosome (EXOSC10 and XRN2) using Cas13d (Fig. S6A and B). This led to 1.7- and 2.1-fold increases in nuclear RNA abundance, respectively, without increasing global Pol II elongation levels (as detected by Pol II phosphoserine 2 staining) or H3K27me3 levels (Fig. 4A; Fig. S6C to H). Under these conditions, chromatin compaction increased by ∼1.25-fold (Fig. 4B), with large DNA-depleted regions visible in the nucleus. This indicates that elevated nuclear RNA levels are sufficient to enhance chromatin compaction independently of transcription and H3K27me3. Consistent with this model, poly(A) RNA FISH revealed nuclear mRNAs within visually apparent interchromatin spaces, while segmentation of RNA-rich domains showed that the most RNA-intense nuclear regions have reduced local DNA association relative to broader RNA-enriched regions and their immediate surroundings (Fig. 4C and D; Fig. S6I and J). These observations indicate that RNA and DNA occupy distinct spatial compartments within the nucleus and support a role for endogenous RNA in chromatin organization.

**Figure 4.**
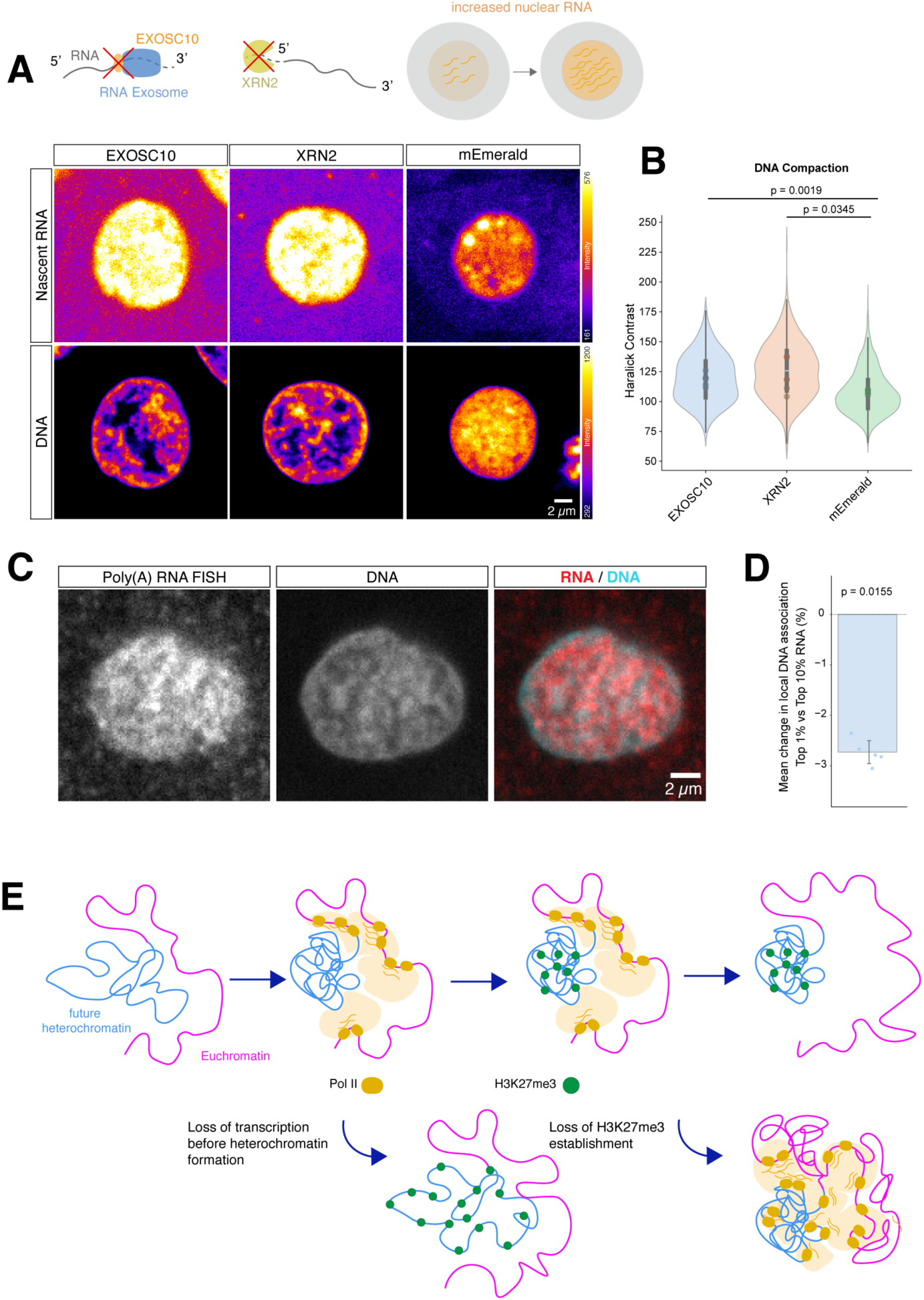
RNA-mediated chromatin compaction. (**A**) Schematic showing the strategy to increase nuclear RNA levels by blocking RNA decay with EXOSC10 or XRN2 knockdown using Cas13d. Metabolic RNA labeling shows significantly increased nuclear RNA levels and significantly increased DNA compaction following EXOSC10 or XRN2 knockdown when compared to a non-targeting control (mEmerald) at 6 hpf. N = 470, 526, and 1,850 nuclei from 6 embryos each for EXOSC10, XRN2, and mEmerald, respectively. Images are single Z-slices of individual nuclei. (**B**) Quantification of DNA compaction from (A). P-values determined by permutation test. (**C**) PolyA RNA FISH shows mRNAs occupying regions of low DNA density at 6 hpf. N = 3,403 nuclei from 6 embryos. (**D**) Quantification of local DNA-depletion at regions with the highest RNA signal from (C) shows that DNA is relatively depleted from sites of high RNA. P-value determined by permutation test. (**E**) Schematic showing the model of RNA-mediated chromatin compaction.

Together, these results demonstrate that RNA is a central determinant of chromatin compaction during early development. Transcription inhibition disrupts chromatin organization despite elevated H3K27me3 and H3K9me3, while increasing nuclear RNA enhances compaction independently of transcription (Fig. 4E). These findings indicate that RNA abundance is a major determinant of the mesoscale chromatin organization during early development.

## Discussion

Here, we show that nuclear organization emerges in an RNA-dependent manner during early development. Using in vivo nanoscale imaging, we find that chromatin compaction arises independently of the PRC2-deposited repressive mark H3K27me3 and is not sufficient to drive transcriptional repression. Although H3K27me3 accumulates at compacted chromatin domains, perturbation of PRC2 activity does not lead to broad chromatin decompaction and instead coincides with target-gene derepression. These results demonstrate that physical chromatin organization, histone modification, and transcriptional repression can be uncoupled in vivo, and identify nuclear RNA as a key determinant of chromatin compaction during early development. Together, our results support a model in which RNA production and accumulation drive the initial physical organization of chromatin, with H3K27me3 functioning within this RNA-dependent framework rather than serving as its primary determinant.

The improved ChromExM workflow developed here enabled nanoscale visualization of the emergence of chromatin organization and its relationship to transcription and histone modification in vivo. Direct visualization revealed that H3K27me3 is associated with, but not required for, mesoscale chromatin compaction. These observations argue that chromatin compaction and PRC2-mediated repression are not causally linked during early development, despite their frequent co-occurrence. This interpretation is consistent with recent work suggesting that chromatin compaction and transcriptional repression can be uncoupled (*29*), and provides direct in vivo evidence that these processes are separable during development. While our results do not entirely rule out the possibility that canonical PRC1, a Polycomb complex capable of directly compacting chromatin (*49*), contributes to chromatin compaction independently of PRC2 during these stages, previous work suggests that non-canonical PRC1 is the predominant PRC1 complex in early zebrafish embryos (*50*). Because non-canonical PRC1 is thought to function primarily via H2A monoubiquitination rather than direct chromatin compaction (*50*, *51*), this suggests that canonical PRC1-mediated compaction is unlikely to be the predominant mechanism underlying the global chromatin organization observed here.

Unexpectedly, our results show that nuclear RNA regulates chromatin organization during early development. Rather than emerging as a downstream consequence of heterochromatin formation, chromatin compaction can arise through RNA-dependent mechanisms even when canonical repressive pathways are disrupted. Together with the observation that elevated H3K27me3 and H3K9me3 are insufficient to establish compaction following transcriptional inhibition, these findings place RNA upstream of chromatin organization and suggest that RNA abundance is a major determinant of chromatin architecture as it first emerges in vivo.

Notably, these findings differ from chromosome conformation capture-based studies reporting that transcription inhibition has relatively modest effects on genome architecture (*7–9*, *13*). While Hi-C measures population-averaged contact frequencies and long-range interaction patterns, ChromExM directly resolves chromatin density and spatial organization within individual nuclei. Thus, transcription-dependent RNA production appears dispensable for many features of genome folding detected by Hi-C, yet critical for the emergence of mesoscale chromatin organization. More broadly, our results support a model in which nuclear RNA acts as a physical organizer of chromatin, within which sequence-specific contacts may further organize. Consistent with this view, RNA occupies interchromatin regions and increasing nuclear RNA abundance is sufficient to enhance chromatin compaction.

These findings raise the possibility that RNA-dependent chromatin organization represents a general principle of nuclear architecture. While we observe this phenomenon operating across whole nuclei during embryonic development, there is precedent for RNA-mediated chromatin compaction at specific genomic loci. For example, XIST RNA promotes compaction of the inactive X chromosome and can precede transcriptional silencing (*52*, *53*). More broadly, diverse coding and non-coding RNAs have been implicated in the formation and organization of nuclear compartments, and in transcriptional regulation (*54–59*). Within this framework, RNA may act not only as a regulator of gene expression but also as a structural component of the nuclear environment capable of shaping chromatin organization. Although the molecular mechanisms remain to be determined, our findings suggest that the accumulation of RNA during development may help organize chromatin into higher-order structures as nuclear architecture first emerges.

In summary, our results identify nuclear RNA as a key determinant of chromatin compaction during early development and demonstrate that chromatin organization, histone modification, and transcriptional repression can be uncoupled in vivo. Although these studies were performed in the uniquely permissive chromatin environment of the early embryo, our findings suggest that RNA is not simply a product of genome activity but may also function as a fundamental organizer of nuclear architecture.

## Supporting information

DataS1

DataS2

DataS3

## Acknowledgments

Imaging was performed at the UCSF Center for Advanced Microscopy using instruments funded in part through an S10 Shared Instrumentation grant (1S10OD017993-01A1). Sequencing was performed at the UCSF Center for Advanced Technology, supported by UCSF PBBR, RRP IMIA, and NIH 1S10OD028511-01 grants. We thank Chris Jeans from the QB3 MacroLab for purifying Rfx-Cas13d protein; and Vijay Ramani, Barbara Panning, and Rebecca Starble for helpful discussions and feedback on the manuscript.

## Declaration of generative AI and AI-assisted technologies in the manuscript preparation process

During the preparation of this work, the authors used chatGPT (OpenAI) for coding aid. The authors reviewed and edited the output as needed and take full responsibility for the content of the published article.

## Author contributions

Conceptualization: MEP

Methodology: AA, NRL, MEP

Investigation: AA, NRL, MEP

Formal Analysis: AA, NRL, MEP

Funding acquisition: MEP

Supervision: MEP

Writing – original draft: MEP

Writing – review & editing: AA, NRL, MEP

## Data, code, and materials availability

Sequencing data generated in this study have been deposited in the NCBI Sequence Read Archive (SRA) under BioProject accession PRJNA1512465. Code used for image analysis and data processing is publicly available at https://github.com/Pownall-lab/zebrafish_image_analysis and will be archived in Zenodo. Other data and materials generated in this study will be made available from the corresponding author upon reasonable request.

## Materials and Methods

### Zebrafish embryo production

Wild-type zebrafish embryos were obtained by randomly selecting and mating adult zebrafish of the TUAB and TL strains, originally obtained from the Zebrafish International Resource Center. Zebrafish were maintained and used under a protocol approved by the UCSF IACUC. Zebrafish adults and embryos were maintained at 28°C. For all experiments, embryos were collected from multiple parents and adults were allowed to mate for no more than 15 minutes to ensure embryos within each clutch developed synchronously.

### Embryo injections

Embryos were injected with 1 nL of nuclease-free water containing various reagents at the one-cell stage. All injections were performed on embryos dechorionated with ∼1 mg/mL Pronase (Sigma-Aldrich P5147) at the one-cell stage.

### Cas13d

Rfx-Cas13d protein was purified from pET-28b-RfxCas13d-His (Addgene Plasmid #141322) by the QB3 MacroLab as previously described (*35*) and stored at -80°C in single-use aliquots with a concentration of 4.6 mg/mL. gRNAs were designed with cas13design (https://cas13design.nygenome.org/) (*60*, *61*). For each target transcript (EZH2, ENSDART00000024722.10; EED, ENSDART00000162928.1; EXOSC10, ENSDART00000137410.3; XRN2, ENSDART00000163071.2), 3 high-scoring gRNAs were selected spanning the coding sequences (Data S1). gRNAs were appended with the Cas13 scaffold and T7 promoter for in-house synthesis by fill-in-PCR and in vitro transcription (IVT) as previously described (*35*) using the HiScribe T7 Quick High Yield RNA Synthesis Kit (NEB, cat no. E2050S) (with overnight transcription reaction). For each target, equimolar amounts of PCR product for each gRNA were pooled prior to IVT. gRNAs were purified by sodium acetate precipitation and stored at -80°C in single-use aliquots. We used gRNAs targeting either mCherry or mEmerald as non-targeting controls; mCherry gRNAs were synthesized by Synthego and mEmerald was synthesized in-house. gRNA sequences are provided in Table S1. Injections were performed by mixing Cas13d protein (3000 ng/µl) and gRNAs (1000 ng/µl gRNA pool) and injecting embryos with 1 nL of the mix directly into the cell at the one-cell stage.

### Pharmacological inhibitors

To inhibit transcription, embryos were either injected with 0.2 ng α-amanitin (Sigma-Aldrich, catalog no. A2263) at the one-cell stage or bathed in 10 μM flavopiridol (Sigma-Aldrich, catalog no. F3055) beginning at the one-cell stage until collection at the desired stage. To inhibit EZH2, embryos were bathed in 25μM tazemetostat (SelleckChem, catalog no. S7128) from the one-cell stage until collection at the desired stage.

### Improved ChromExM

ChromExM was performed as previously described (*33*) with the following modifications. Embryos were fixed directly without manual deyolking. Embryos were transferred into fixation solution consisting of 3% paraformaldehyde (PFA; Electron Microscopy Sciences, catalog no. 50-980-487) and 0.1% glutaraldehyde (GA; Electron Microscopy Sciences, catalog no. 50-262-10) in 1x PBS in a low retention round bottom tube (Eppendorf catalog no. 022431102) and fixed for 15 min at room temperature (RT) with rotation. Fixation solution was prepared fresh for each experiment and used within 1 hr. Embryos were washed twice briefly in PBS and immediately post-fixed (also referred to as anchoring) in 20% acrylamide (AAm; Sigma-Aldrich, catalog no. 01697-500ML) and 4% PFA at 4°C with rotation overnight (up to ∼18 h) to enable covalent anchoring of proteins to the hydrogel while limiting excessive intermolecular crosslinking.

Following post-fixation, embryos were washed twice with PBS and yolks were disrupted by brief vortexing (5-10 s intervals, typically 1-3 rounds of vortexing) to release the yolks. Embryos were washed once more with PBS, then transferred to a new tube containing fresh PBS to remove residual yolk and could be stored at 4°C for up to ∼1 month prior to gelation.

The first gelling solution consists of 10% acrylamide, 0.1% N,N’-(1,2-dihydroxyethylene)bisacrylamide (DHEBA; Millipore-Sigma, catalog no. 294381-5G), and 19% sodium acrylate (SA; Pfalz and Bauer, catalog no. S03880-25G). Sodium acrylate was prepared as a 38% stock solution immediately before use and centrifuged at 4,000 rpm for 5 min to remove impurities, and each lot was assessed for quality as previously described (*62*, *63*). This solution can be prepared in advance and stored at 4°C. At the time of gelation, the first gelling solution was activated by the addition of ammonium persulfate (APS; Sigma-Aldrich catalog no. A3678-25G) and N,N,N’,N’-tetramethylethylenediamine (TEMED; Thermo Scientific, catalog no. 17919) to a final concentration of 0.05% each immediately before use. Embryos were incubated in activated first gelling solution for 5-15 min on ice. Then, one sample at a time, embryos were transferred to a round #1.5 12 mm coverslip on a gelation chamber assembled from two stacks of #2 22 mm and one #0 22 mm square coverslips superglued onto a glass slide on either side of the 12 mm coverslip. Excess solution was wicked away from the embryos on the coverslip using a kimwipe and 85 µL of freshly activated first gelling solution was immediately added to the embryos. Embryos were manually positioned on the coverslip using forceps to ensure spacing and contact with the glass, and the gelation chamber sealed with a 22 mm coverslip to achieve an initial gel thickness of ∼370 µm. Chambers were transferred to a humidified gas exchange chamber, purged with nitrogen for 10 minutes, and incubated at 37°C for 1.5 h.

Following gelation, individual embryos were isolated from the polymerized gel using a 1 mm tissue biopsy punch. Gel disks containing embryos were immediately transferred into denaturation buffer (200 mM NaCl, 50 mM Tris, 200 mM sodium dodecyl sulfate, pH 6.8) and incubated at 82°C (temperature at the sample) for 3 hrs with shaking (typically 450 rpm). Gels were then washed extensively in PBS (3-5 washes, 5-10 min each) and transferred to ultrapure water for expansion (two 30 min washes followed by one overnight wash).

After the first expansion, embryos were re-embedded in a neutral hydrogel consisting of 10% acrylamide and 0.05% DHEBA in ultrapure water. To activate polymerization, APS and TEMED were added to a final concentration of 0.05%. Embryos were washed in this activated neutral gelling solution three times for 20 min on a rocker, using freshly activated gelling solution for each wash. After the final wash, the gels were sandwiched between two no.1.5 coverslips after removing excess solution, transferred to a humidified gas exchange chamber, and purged with nitrogen for 7 min. Gelation then proceeded for two hours at 37°C. After gelation was complete, the gels were retrieved and stored at room temperature in PBS overnight. Subsequently, embryos were embedded in a second swellable hydrogel consisting of 10% acrylamide, 19% SA, and 0.1% N,N′-methylenebis(acrylamide) (BIS; Alfa Aesar, catalog no. J66710-14) in PBS. The gelling solution was activated by the addition of 0.05% APS and TEMED. Embryos were washed three times for 15 minutes on ice on a rocker in freshly activated gelling solution. After the final wash, the gels were sandwiched between two no.1.5 coverslips after removing excess solution, transferred to a humidified gas exchange chamber, and purged with nitrogen for 7 min. Gelation then proceeded for two hours at 37°C

Following the final gel polymerization, crosslinks from the first and second hydrogels were hydrolyzed by incubating the gels in 200 mM NaOH for 2 hrs at RT with rocking. Gels were then neutralized by washing 2-3 times for 30 min with excess PBS. During the final PBS wash, gels were trimmed to isolate only the embryo in order to reduce volume and improve subsequent staining efficiency.

After staining (see below), gels were expanded in ultrapure water by exchanging water once per hour for two hours, followed by full expansion in fresh ultrapure water for at least 6 h in the dark.

Gels were sliced manually along the Z-axis with a fresh razor blade and mounted in no. 1.5 glass-bottom MatTek dishes (MatTek, catalog no. P50G-1.5-30-F) coated with Poly-L-lysine (Sigma-Aldrich, catalog no. P8920). Typically, 1-2 embryos were mounted per dish. Approximately 25 µL of 10 µM SYTOX Green (Invitrogen, catolog no. S7020) was added directly to each gel slice in the dish prior to sealing. A #1.5 22 × 22 mm coverslip (Electron Microscopy Sciences, catalog no. 7220401) was placed on top of the gels and sealed using Z-dupe (Henry Schein, catalog no. 1026526). Samples were stored at 4°C protected from light and imaged within 3-4 days.

### ChromExM DNA labelling

We previously observed extensive photobleaching of SYTOX Green when used for DNA labeling (*33*). This strategy led to a 64% reduction in signal over 300 frames (acquired as a 2D timeseries of the same Z-plane) (Fig. S1D). We reasoned that SYTOX Green may be undergo less photobleaching if the dye is present in excess in the gel at the time of imaging, which could enable exchange of bleached dye molecules. To achieve this, we both increased the amount of dye used for pre-expansion staining (5 µM in HBSS (Hanks’ Balanced Salt Solution; Thermo Fisher Scientific, catalog no. 14170-112) for 30 minutes before the final expansion) and added SYTOX Green (10 µM in H_2_O) directly to the gels at the time of mounting and measured its bleaching (34 % reduction in mean intensity over 300 frames; Fig. S1E). We measured bleaching during each experiment and applied a correction factor on a per-slice basis using the measured bleaching curve to account for the remaining bleaching that occurs during imaging.

### Development of improved ChromExM

We sought to overcome two major bottlenecks in ChromExM. First, because samples cannot be stored after fixation before gelation (*33*), the first day of the experiment can be excessively long (>20 hrs) (Fig. S1A). To overcome this, we tested several different fixation and anchoring (also referred to as post-fix) strategies with overnight incubation or overnight storage. We found that fixation in 3% PFA and 0.1% GA in 1x PBS worked well. Using an overnight post-fix (4% PFA, 20% AAm in 1x PBS) (*64*, *65*), we observed that with increased denaturation temperature and time, cells appeared intact with minimal defects (Fig. S1C). The second challenge is that not all epitopes, such as H3K27me3, are well-detected with our standard post-expansion staining approach (*33*). This could be due to 1) epitope masking during fixation; 2) steric hinderance caused by the acryloyl anchoring step; or 3) epitope or signal loss during expansion. We have excluded the possibilities of (1) and (2) by performing primary antibody staining after embryos are fixed and anchored following our newly established protocol but prior to gelation. Secondary antibody staining is performed after gel polymerization to avoid bleaching of the fluorophores.

### ChromExM NHS-ester staining

NHS-ester staining was performed as previously described (*63*). Briefly, just prior to the final expansion in water, gels were stained in 20 µg/mL Abberior STAR 635 NHS ester (Sigma-Aldrich, catalog no. 30558) in 100mM NaHCO_3_ for 1.5 hrs at RT on a rocker. Gels were then washed 3×20 min with PBST (1x PBS with 0.1% Tween-20) and expanded as described above.

### Once expanded ChromExM

Embryos were fixed and anchored as described above. Before gelation, embryos were blocked in 10% bovine serum albumin (BSA; Sigma-Aldrich A9647-100G) in PBS containing 0.5% Triton-X-100 (PBSTr) for 2 hrs at RT and incubated with primary antibody (H3K27me3, Active Motif 39155, 1:25) in 10% BSA in PBSTr overnight at 4°C. Samples were washed 3×10 min with PBSTr, then subjected to an additional anchoring step by incubation in 1% PFA, 2% AAm in 1xPBS at 4°C overnight. Then the first gelation and denaturation proceeded as described above. After extensive PBS washing, samples were stained with secondary antibody (Goat anti-Rabbit IgG Alexa Fluor 568, Invitrogen A11036, 1:100) in 2% BSA in PBST for ∼48 hrs at 4°C. Samples were washed 3×10 min in PBST at RT, then stained with tertiary antibody (Donkey anti-Goat Alexa Fluor 568, Invitrogen, A11057, 1:100) in 2% BSA in PBST for ∼48 hrs at 4°C. Samples were washed 3×10 min in PBST at RT, stained with SYTOX Green (5 µM in HBSS for 30 min at RT), washed with PBST, and expanded in ultrapure water. Samples were then mounted as described above. H3K27me3 was visualized with this once-expanded approach for optimal signal.

### Unexpanded fixed imaging

Staining was carried out as previously described with modifications (*33*). Embryos were fixed in 4% PFA in 1x PBS at 4°C overnight, washed with PBSTr, then dehydrated in a series of methanol/PBSTr washes (25%, 50%, 75%, 100% methanol) and stored at -20°C for at least 2 hours. Embryos were rehydrated with a reverse series of methanol/PBSTr washes, incubated for 20 min on ice in ice-cold 100% acetone, washed twice for 5 min with PBSTr, then permeabilized with 10 µg/mL Proteinase K (Millipore 70663-4) in PBSTr for exactly 2 minutes. Embryos were then washed twice with PBSTr and fixed with 4% PFA at RT for 30 min. After fixation, embryos were washed twice for 10 minutes with PBSTr, then blocked for 2-3 hrs at RT in 10% BSA in PBSTr. Then, embryos were incubated at 4°C overnight with primary antibody (H3K27me3, Active Motif 39155, 1:500; H3K9me3, Abcam ab8898, 1:500; Pol II pSer2, Abcam ab193468, 1:500) in 10% BSA in PBSTr, washed with PBSTr, and incubated in secondary antibody (Goat anti-Rabbit IgG Alexa Fluor 568, Invitrogen A11036; Goat anti-Rabbit IgG Alexa Fluor Plus 647, Thermo Fisher Scientific A32733; Goat anti-Rabbit IgG Alexa Fluor Plus 488, Thermo Fisher Scientific A32731; all 1:1000 dilution) in 10% BSA in PBStr for two hours at RT. Embryos were then washed 3×10 minutes with PBSTr, incubated with 5 µg/mL DAPI (Invitrogen, catolog no. D1306) in PBSTr for 18 minutes at RT, washed twice more with PBSTr, then stored at 4°C for up to 3 days. Embryos were mounted in 0.8% low-melting point agarose in PBS for imaging.

### Unexpanded RNA SABER-FISH

RNA SABER-FISH was performed as previously described (*66*) with modifications. We used a Locked Nuclei Acid (LNA) polyT probe (+T+T+TTT+TTT+TTT+TTT+TTT+T+T+TTTTACATCATCAT; IDT) to detect polyA mRNAs (*67*, *68*). The primary probe was extended using Primer Exchange Reaction (PER). Briefly, PER reactions were assembled in 1x PBS containing MgSO_4_, dATP, dCTP, and dTTP, together with clean.G oligo, Bst LF polymerase (NEB, catalog no. M0275L), PER hairpin (ACATCATCATGGGCCTTTTGGCCCATGATGATGTATGATGATGTTTTTTT), and primary probe template (*66*). Reactions were incubated sequentially at 37°C to allow concatemer extension, heat-inactivated at 80°C for 20 min, purified by sodium acetate/ethanol precipitation, and resuspended in 30 µL nuclease-free water. Probe length was verified by agarose gel electrophoresis prior to use and probes were approximately 500 nucleotides in length.

Embryos were fixed in 4% PFA in 1x PBS at 4°C overnight, washed with PBSTr, and optionally stored at 4°C. Embryos were then washed in 2x SSCT (2x SSC, 0.5% Tween-20) before equilibration with two 15 min washes in pre-warmed hybridization wash buffer (40% formamide, 2x SSC, 0.5% Tween-20) at 37°C. Purified PER-extended probes were diluted 1:100 in hybridization buffer (50% formamide, 2x SSC, 0.5% Tween-20, 10% dextran sulfate (Sigma-Aldrich, catalog no. D8906-50G)), denatured at 60°C for 3 min, then hybridized with embryos overnight at 37°C with shaking at 450 rpm. Embryos were subsequently washed twice in pre-warmed hybridization wash buffer at 60°C, followed by washes with 2x SSCT at 37°C, then with PBS at room temperature.

For detection, fluorescent imager oligos (5ATTO647NN/ttATGATGATGTATGATGATGT/3InvdT) were diluted to 1 µM in pre-warmed 1x PBS and hybridized with embryos for 2-3 h at 37°C protected from light. Embryos were then washed extensively in pre-warmed PBSTr at 37°C, stained with DAPI, and stored protected from light at 4°C prior to imaging.

### Nascent RNA imaging

Embryos were injected with 20 mM 5-Ethynyl Uridine (5-EU; Vector Laboratories, catalog no. CCT-1261-25) at the one-cell stage. EU was detected with 5 µM AZDye 647 Picolyl Azide (Vector Laboratories, catalog no. CCT-1300-1) using the Click-&-Go Cell Reaction Buffer Kit (Vector Laboratories, catalog no. CCT-1263) as previously described (*33*).

### Spinning Disk Confocal microscopy

Images were acquired using a CSU-W1 Spinning Disk microscope on a Nikon Ti body equipped with a Zyla sCMOS 4.2 megapixel camera. Unexpanded images were acquired with a Nikon 60x 1.2 NA water-immersion objective. ChromExM images were acquired with a Nikon 10x 0.45 NA air or a Nikon 40x 1.15 NA water-immersion objective. Images were acquired with the following voxel sizes (X, Y, Z µm): unexpanded and once-expanded ChromExM (0.109, 0.109, 0.3); ChromExM, 40x (0.163, 0.163, 0.5); ChromExM 10x (0.65, 0.65, 5). Stacks were acquired in the order of channel, slice such that the entire Z-stack was imaged in one channel before imaging the next channel. Channels were acquired from longest wavelength to shortest. We used 405 nm (100 mW), 488 nm (150 mW), 561 nm (100 mW), and 639 nm (160 mW) laser lines with 447/60m, 525/50m, 607/36m, and 685/40m emission filters, respectively. The microscope was controlled by Micromanager (*69*, *70*).

### Image analysis

Image analysis was performed using custom written python scripts available at https://github.com/Pownall-lab/zebrafish_image_analysis. Analyses utilized functionality from NumPy, SciPy, pandas, scikit-image, PyTorch, and OpenCV (*71–76*).

#### Unexpanded image processing

We implemented a fully automated pipeline for processing unexpanded images of embryos to segment nuclei, quantify nuclear intensity features, and perform downstream statistical analysis and plotting without manual intervention. A JSON configuration file specifies inputs, outputs, channel mapping, and all processing parameters, enabling reproducible and standardized analysis across experiments. Images were read from the native MicroManager (*69*, *70*) directory structure to generate maximum intensity Z-projections and single-channel 3D image stacks. Maximum projections were used for visual inspection, while all quantitative analyses were performed on 3D images. Single-channel images were optionally subjected to primary background subtraction using an acquired “dark” image to correct for camera offset.

Nuclei were segmented in 3D from the DNA channel using a custom Cellpose-SAM model fine-tuned on our data (see Cellpose training). Border labels were excluded and segmentation masks were filtered based on size and sphericity (typically >25,000 voxels and >0.55 sphericity) to remove small background objects and mitotic nuclei.

To reduce residual non-specific signal, we implemented a secondary background subtraction step in which cytoplasmic regions were automatically identified using nuclear masks. Background intensity was estimated per image and aggregated across all embryos within an experiment to compute a global background value for each channel. This global per-channel background was then subtracted from all images.

Intensity features were computed from background-subtracted single-channel images for each nucleus using the corresponding segmentation masks. Haralick contrast (*77*) was calculated on a per-slice basis in 2D using a masked gray-level co-occurrence matrix restricted to pixels within each nucleus, and values were averaged across Z-slices. This served as a proxy for DNA compaction in diffraction-limited images by measuring the variability of DNA signal in nuclei (*46*, *78*). Contrast was computed using a pixel distance of 2, averaged across multiple angles.

#### Cellpose training

To segment nuclei in unexpanded images, we trained a fine-tuned Cellpose-SAM (CPSAM) model using targeted 2D slices extracted from representative 3D image stacks. Training data were generated from input stacks spanning multiple developmental stages and experimental perturbations, ensuring representation of both typical and challenging nuclear morphologies, with slices sampled across XY, XZ, and YZ planes. Initial 2D masks were generated using the pretrained CPSAM model and manually curated to remove or correct poorly segmented examples. A subset of image-mask pairs (∼5–10%) was withheld for testing. The model was trained using Cellpose with a learning rate of 1×10⁻⁵, weight decay of 0.1, and 100 epochs on GPU. For segmentation of 3D images, the trained model was applied using Cellpose’s 3D segmentation mode.

#### ChromExM image processing

We developed an automated pipeline for segmentation and quality control of ChromExM images. A JSON configuration file specifies inputs, outputs, channel mapping, and processing parameters. The pipeline performs channel splitting, optional primary background subtraction using a dark image, and nucleus segmentation.

Nuclei were segmented from the DNA channel using a custom 2.5D U-Net inference pipeline (see ChromExM nucleus segmentation model and training section), applied slice-by-slice to generate 3D nuclear masks. Segmentation outputs were reconstructed into 3D labeled volumes for downstream analysis.

Segmentation quality was assessed using automated metrics, including mask coverage fraction, border-touching fraction, and slice-wise solidity, which were used to identify potential segmentation errors for manual review.

#### Extended z-stack stitching

For nuclei that exceeded the axial imaging range of the piezo Z-stage on our microscope, two sequential 3D stacks were acquired from the same field using the piezo for each stack and then stitched computationally to generate an extended z-stack. Stitching was performed using a custom Python script that determined the z-overlap by anchoring to the final slice of the first stack and searching for the best-matching slice within the beginning of the second stack. Matching was performed on the DNA channel after simple background subtraction using normalized cross-correlation and a gradient-magnitude transform of the DNA channel. The overlap length was defined by the matched slice in the second stack, and the stitched volume was generated by retaining the full first stack and appending only the non-overlapping portion of the second stack. Datasets for which a reliable overlap could not be identified were excluded from downstream analysis.

To correct for small rigid XY drift between sequential acquisitions, a single global integer translation was estimated using phase correlation and applied uniformly to the second stack prior to stitching. The translation was computed using the first retained slice of the second stack to ensure accurate alignment at the stitching boundary.

#### ChromExM nucleus segmentation model and training

For ChromExM nucleus segmentation, we trained a 2.5D U-Net-based convolutional neural network in PyTorch. The model took three adjacent DNA slices as input channels and predicted a single-channel nucleus probability map for the center slice. Architecturally, the network consisted of an encoder-decoder U-Net with residual convolutional blocks using GroupNorm and an atrous spatial pyramid pooling (ASPP) bottleneck. Training data consisted of annotated 3D volumes spanning multiple developmental stages and experimental conditions. Binary masks were generated using the previously published ChromExM segmentation approach (*33*) and from prior model iterations, and were manually curated to correct segmentation errors. Volumes were split into training and validation sets at the volume level (80/20 split). Each volume was normalized by percentile clipping (1st-99th percentile) and divided into overlapping 512×512 pixel patches with 50% stride. In total, 16 annotated volumes yielded approximately 80,096 training patches and 30,462 validation patches. Patch sampling was biased to retain interior and boundary-containing nuclear regions while downsampling background-only patches. On-the-fly augmentation included random horizontal and vertical flips, 90° rotations, elastic deformations, intensity scaling and offset, Gaussian blur, and Gaussian noise. The network was optimized using Adam (learning rate 1×10^-4^; batch size 12) with a combined Focal Tversky and binary cross-entropy loss, and the model with the lowest validation loss over 50 epochs was selected for downstream use. Model performance was monitored using validation loss and Dice coefficient (threshold 0.5) during training.

For inference, 3D nucleus masks were generated from full image volumes using a sliding-window prediction strategy. Each volume was normalized by percentile clipping (1st-99th percentile), and predictions were computed slice-by-slice using three adjacent slices as input channels. Overlapping tiles were combined using Hann-window weighting to minimize edge artifacts. A threshold (default 0.5) was applied to probability maps to produce binary masks, followed by 3D connected-components labeling. Only the largest connected component was retained to remove small speckles and background fragments.

#### Photobleaching measurement

Photobleaching of SYTOX Green was quantified from time-series ChromExM imaging of individual nuclei. Single-slice time series (300 frames) were acquired and treated as 3D stacks with time encoded along the Z-axis. For each dataset, corresponding nuclear segmentation masks were applied on a per-frame basis, and mean fluorescence intensity within the nuclear mask was computed independently for each frame to generate a time-resolved intensity trace. For each experiment, traces from >9 nuclei were aggregated, and the mean intensity profile across nuclei was used to estimate the bleaching curve.

#### Bleach correction

Photobleaching correction was applied to ChromExM image stacks using an experimentally derived bleaching profile (see Photobleaching measurement). Briefly, bleaching was quantified from a representative set of nuclei imaged under identical acquisition conditions for each experiment, and the resulting average intensity profile across acquisition frames was used to estimate signal loss due to photobleaching. A smooth bleaching curve was fit to these measurements and used to derive correction factors for each acquisition frame, which were applied to all image stacks from the corresponding experiment. Corrected intensities were clipped to the uint16 range without additional rescaling. Bleach-corrected image stacks were used for all subsequent visualization and quantitative analyses.

#### Radial autocorrelation

Radial chromatin autocorrelation was computed from ChromExM DNA images using a custom Python script. For each nucleus, the DNA image volume was normalized within the nuclear mask by percentile clipping (1st-99th percentile). Autocorrelation was then computed independently for each Z-slice using a fast Fourier transform (FFT) based approach after subtraction of the mean nuclear intensity. The autocorrelation of the masked signal was normalized by the autocorrelation of the corresponding nuclear mask to correct for boundary effects. The resulting 2D autocorrelation maps were radially averaged to generate slice-wise autocorrelation profiles, which were normalized such that the autocorrelation at a distance of 0 was equal to 1 (i.e., C(0)=1). Slice-wise radial profiles were then combined into a single per-nucleus autocorrelation curve using a weighted average at each radius based on the number of contributing pixel pairs.

#### Chromatin occupancy

Local chromatin occupancy was computed from ChromExM DNA images using a sliding 3D window approach. DNA intensities were first normalized within the nuclear mask by percentile clipping, and voxels with normalized intensity values greater than 0.1 were classified as chromatin. For each voxel within the nuclear mask, local chromatin occupancy was calculated as the fraction of neighboring voxels classified as chromatin within a 3D window of 2×6×6 voxels (Z, Y, X). This produced a voxel-wise map of local chromatin occupancy within each nucleus.

#### Expansion factor measurement

Expansion factor was determined based on nuclear cross-sectional area as previously described (*33*). For expanded samples, ChromExM images acquired with a 10x objective were segmented from the DNA channel using an automated 3D pipeline consisting of global Otsu thresholding, slice-wise hole filling, removal of small objects, 3D connected-component labeling, and downstream filtering of border-touching, small, and low-sphericity objects. Unexpanded images were processed as described above. For each segmented nucleus, cross-sectional area was measured from the middle z-slice of the nucleus, defined as the central slice within the z-extent of its segmentation mask. Expansion factor was then computed by comparing nuclear cross-sectional areas between expanded and unexpanded samples. We applied a conservative 18x expansion-factor correction to all expanded images unless otherwise noted such that the scale bar represents the biological scale after accounting for physical expansion.

#### Once-expanded ChromExM expansion factor

To determine the expansion factor of once-expanded samples, embryos were imaged with a 10x objective and processed using the above described pipeline and segmentation model for unexpanded embryos. Nuclear cross-sectional area was then calculated as described above. We applied a 4.5x expansion-factor correction to all once-expanded images.

### qPCR

RNA was extracted by lysing 30 embryos at 6 hpf in 500 µL of Trizol (Invitrogen, catalog no. 15596026) using a hand-held biovortexer. Lysates were stored at -20°C. Subsequently, samples were thawed and 100 µL chloroform was added to the lysate and mixed thoroughly. The samples were centrifugated at 1200 x g for 15 min at 4°C. Clear supernatant was transferred to a fresh tube to which 250 µL of isopropanol was added, incubated for 10 min on ice, and centrifugated at 1200 x g for 15 min at 4°C to pellet the RNA. Supernatant was removed and the pellet was washed twice with 500 mL ice cold 75% ethanol by briefly vortexing and then centrifugating at 7500 x g for 5 min at 4°C. The RNA pellet was air-dried and reconstituted in 44 µL nuclease-free water. Subsequently, RNA was treated with 1 µl TURBO DNase (Thermo Fisher, catalog no. AM2238) in TURBO DNase buffer for 30 min at 37°C and then purified using the Monarch Spin RNA cleanup kit (NEB, catalog no. T2110) according to the manufacturer’s protocol. cDNA was synthesized using the LunaScript RT SuperMix Kit (NEB, catalog no. E3010) and qPCR was performed using the Luna Universal qPCR Master Mix (NEB, catalog no. M3003) on a QuantStudio 5 Real-Time PCR Instrument (Applied Biosystems) using the SYBR/FAM setup. All primers used for qPCR had an efficiency >100%, except for XRN2, which was 80%. Primers sequences are available in Data S2. qPCR data were analyzed using the ΔΔCt method, normalized to β-actin expression (*79*), and presented as fold change in expression relative to control samples. Relative expression was calculated as 2^^(-ΔΔCt)^, such that the matched control samples were normalized to a value of 1. Data are presented as mean ± SD of biological replicates. Knockdown efficiency was quantified using paired ΔΔCt values by comparing each knockdown sample to its matched sibling control. Each experiment was run with technical triplicates for each of the three biological replicates. Statistical significance was assessed using a two-tailed paired t-test performed on ΔCt values rather than transformed fold-change values.

### SLAM-seq

SLAM-seq was performed as previously described (*40–42*). Briefly, embryos were injected with Cas13d and gRNAs, followed by injection with 60 mM 4sU (Sigma-Aldrich, catalog no. T4509) in a second needle, and reared in the dark until mEmerald-targeted embryos reached shield stage (6 hpf). Then, 30 embryos were collected, lysed in 500 µL Trizol, and stored at -20°C. Samples were thawed, 100 µL chloroform was added, and then spun at 16,000 g at 4°C for 15 minutes. The aqueous phase was transferred to a fresh tube and mixed with 250 µL isopropanol, 2.5 µL fresh 10 mM DTT (Dithiothreitol; Invitrogen, catalog no. P2325), 1 µL 20 mg/mL glycogen (Thermo Scientific, catalog no. R0551), and then spun at 16,000 g at 4°C for 20 minutes. The supernatant was removed and the pellet was resuspended in 500 µL ice-cold 70% ethanol with 0.1 mM DTT before being spun at 7,000 g at RT for 5 minutes. The pellet was air dried and resuspended in nuclease-free water containing 1 mM DTT. RNA (1 µg) was alkylated with in a 50 µL reaction containing 10 mM iodoacetamide (Sigma-Aldrich, catalog no. I1149-5G) in a buffer of 50 mM NaPO_4_ (pH 8; Alfa Aesar, catalog no. J60825.AK) and 50% DMSO by incubating at 50°C for 15 minutes. The reaction was quenched by the addition of 1 µL 1M DTT, then the RNA was precipitated by mixing with 1 µL 20 mg/mL glycogen, 5 µL 3M sodium acetate, and 125 µL 100% ethanol and incubating at -80°C for 30 minutes. The RNA was pelleted at 4°C and washed with ice-cold 75% ethanol, then air dried and resuspended in nuclease free water.

Libraries were prepared from 500 ng of RNA using the QuantSeq 3’ mRNA-Seq V2 Library Prep Kit FWD with UDI (Lexogen; cat no. 191.24) following the manufacturer’s protocol. Libraries were amplified for 13 cycles based on qPCR and their quality and fragment size was assessed using an Agilent 2100 Bioanalyzer. Libraries were pooled and sequenced on a NovaSeqX with SE100 single-end reads. We performed SLAM-seq on two replicates which were injected and collected on separate days. All other steps after lysis were performed together for the replicates.

### SLAM-seq Analysis

Sequencing data were managed with LabxDB seq (*80*) and processed using LabxPipe (https://github.com/vejnar/LabxPipe) and Slamdunk (*40*) (Data S3). Raw sequencing reads were downsampled to 50 million reads per sample using seqtk, adapter trimmed using readknead, and aligned to the Danio rerio GRCz11 reference genome. T>C conversion calling and quantification of labeled transcripts were performed using Slamdunk with a minimum alignment identity of 0.95, 5 nt 5′ trimming, and a minimum variant fraction of 0.2. Multimapping reads were retained with a maximum multimapping threshold of 100.

Differential analysis was performed in R using edgeR (*81*). Slamdunk count tables were mapped to Ensembl gene annotations and collapsed to gene-level counts prior to analysis. Differential analysis was performed on labeled RNA counts using trimmed mean of M-values (TMM) normalization and edgeR generalized linear modeling. Genes with Log2 fold-change > 4 and FDR < 0.05 were considered significantly differentially expressed. Volcano plots and overlap analyses were generated using ggplot2 and dplyr.

### CUT&RUN analysis

Published H3K27me3 CUT&RUN data from 6 hpf embryos (*43*) (NCBI SRA SRP324442) were processed using LabxPipe. Reads were aligned to the Danio rerio GRCz11 reference genome using Bowtie2 (*82*). Duplicate reads were removed using samtools-based deduplication (*83*), and normalized genome-wide coverage tracks (bigWig) were generated using GeneAbacus (https://github.com/vejnar/GeneAbacus). To assess H3K27me3 occupancy at genes transcriptionally activated following PRC2 depletion, H3K27me3 CUT&RUN signal was measured in the set of genes significantly upregulated in both EED- and EZH2-targeted embryos. Shared non-differentially expressed genes identified from both conditions were used as a background set. Metagene profiles were generated using deepTools computeMatrix in scale-regions mode (*84*). H3K27me3 signal was quantified across gene bodies with 2 kb flanking regions upstream of the transcription start site (TSS) and downstream of the transcription end site (TES). Mean normalized H3K27me3 signal was calculated for each gene and compared between gene sets using a one-sided Mann-Whitney U test.

### Statistical analysis

Statistical analyses were performed using custom Python scripts. For analyses of ChromExM datasets, statistical comparisons were performed using measurements from individual nuclei, with nuclei sampled from at least two embryos. For analyses of unexpanded imaging, measurements from individual nuclei were first averaged within each embryo, and per embryo means were used for statistical comparisons.

Group comparisons were performed using two-sided permutation tests. For unexpanded analyses, the difference in group means was used as the test statistic. For ChromExM analyses, including chromatin occupancy and autocorrelation length measurements, the difference in group medians was used as the test statistic. Statistical significance was determined by comparing the measured difference between groups to the distribution of differences obtained by permutation of group labels.

Unless otherwise stated, graphs show the median ± interquartile range, with individual points representing measurements from each nucleus or embryo. The statistical test, sample size, and exact P values are indicated in the corresponding figure legends.

---

**Figure S1.**
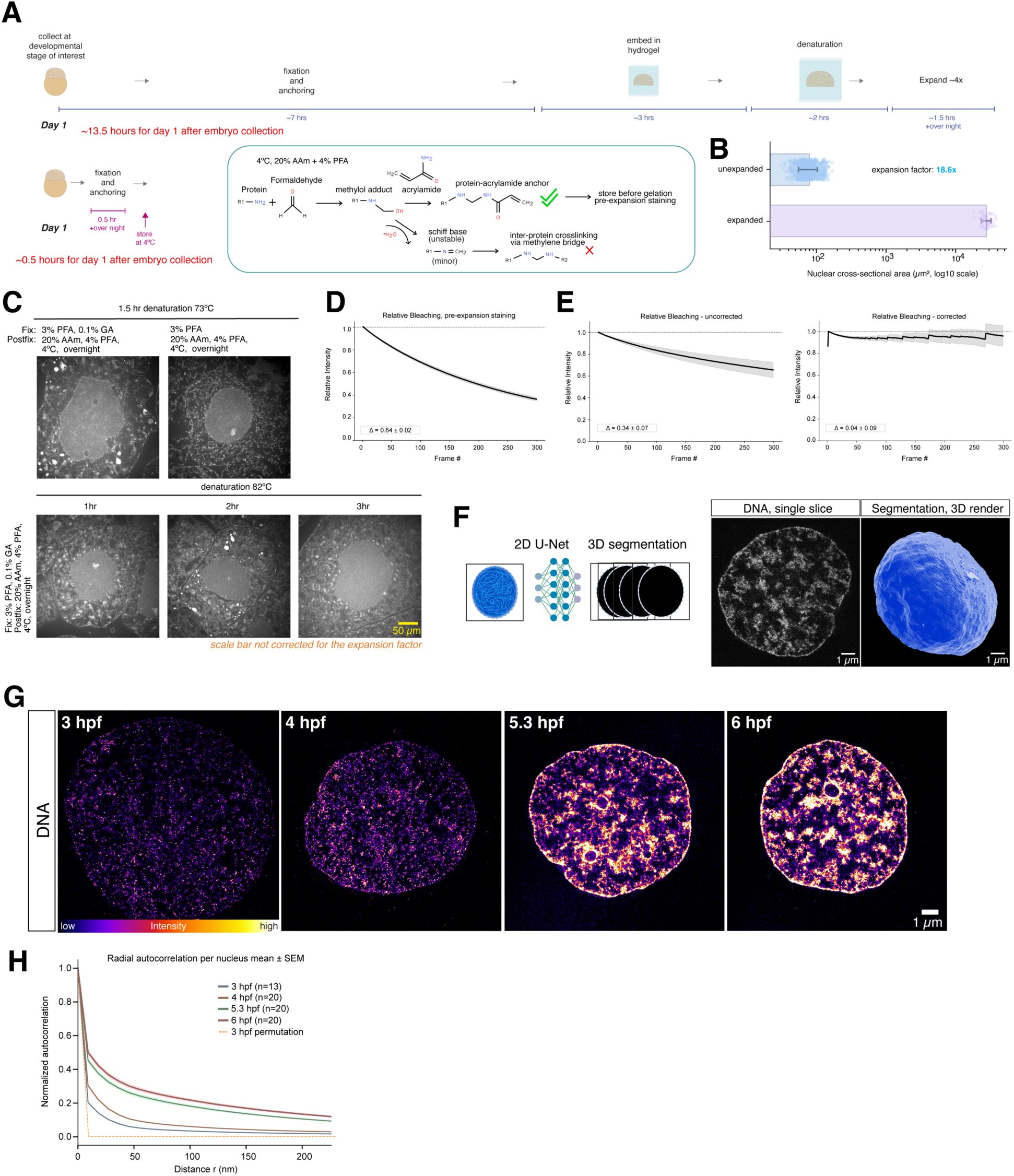
Improvements in ChromExM. (**A**) Schematic of the new ChromExM workflow showing how modified anchoring conditions and solutions support protein-acrylamide anchoring rather than inter-molecular crosslinks. This reduces the active time needed on Day 1 of the experiment from approximately 13.5 hours to 0.5 hours and enables sample storage prior to beginning the expansion process. (**B**) Expansion factor measured by nuclear cross-sectional area (linear expansion factor = 18.6; n = 894 nuclei from 6 embryos and 52 nuclei from 2 embryos, for unexpanded and expanded conditions). (**C**) Representative images of NHS-ester pan staining of proteins showing poor protein retention and ultrastructural preservation (seen has apparent holes and tears throughout the cytoplasm) when samples are stored overnight without using the improved ChromExM protocol. N = 5-9 cell from 2 embryos in each condition. Scale bars are not corrected for the expansion factor. (**D**) Measurements of DNA dye intensity over a 300 frame time series acquisition with pre-expansion staining only (n = 14 nuclei from 1 embryo). (**E**) Measurements of DNA dye intensity over a 300 frame time series acquisition before and after applying bleach correction with pre- and post-expansion staining (n = 18 nuclei from 6 embryos). (**F**) Schematic of the U-Net segmentation strategy and representative segmentation example. (**G**) Images shown in Fig. 1B with display setting matched across all images for visual comparison. (**H**) Radial autocorrelation curves corresponding to the images in (F). As a control, an intensity permutation from 3 hpf images shows all detected autocorrelations are nonrandom.

**Figure S2.**
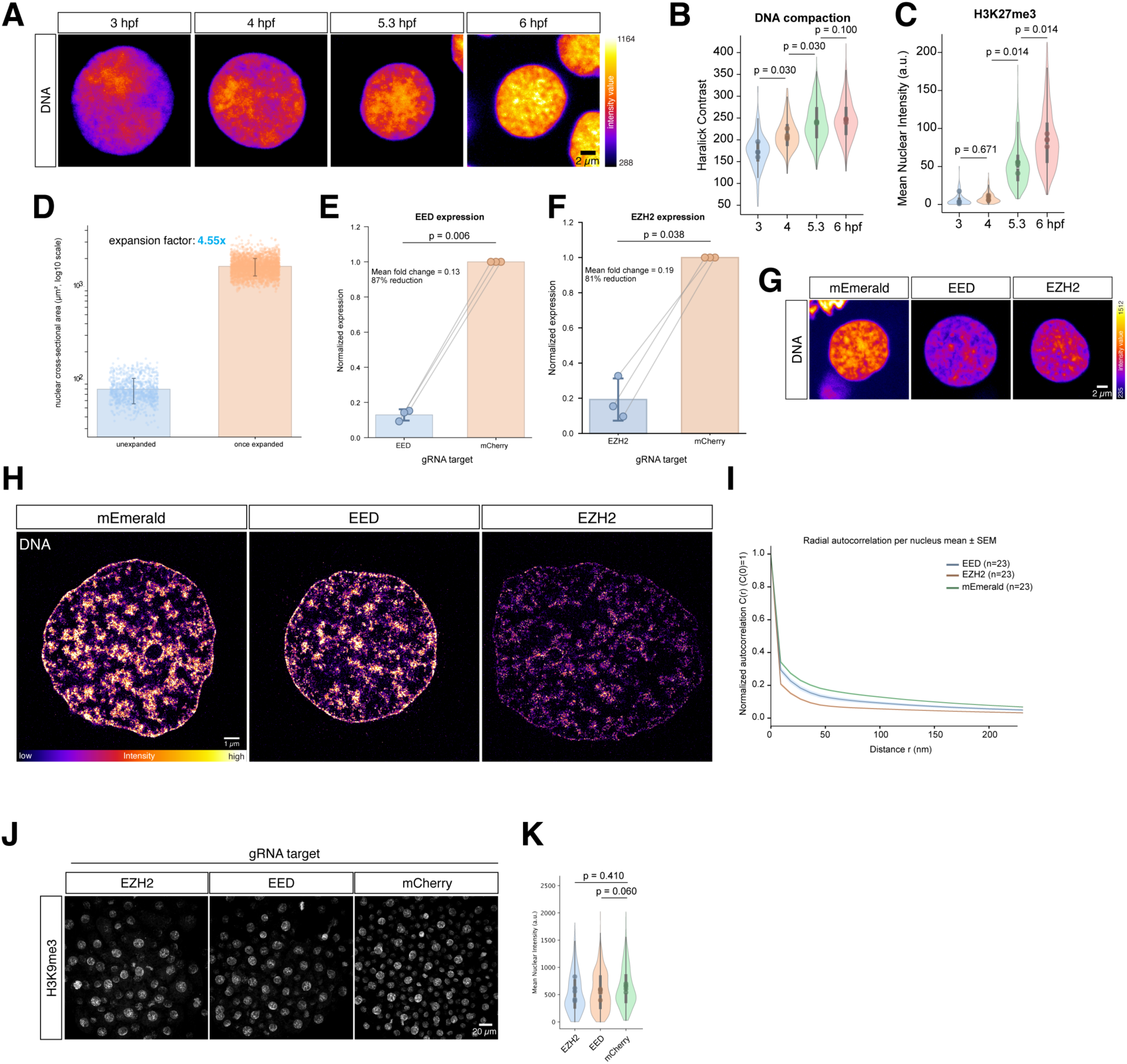
Additional characterization of H3K27me3 and PRC2 knockdown. (**A**) Images of DNA corresponding to Fig. 2A. Images are maximum intensity projections of individual nuclei. N = 103, 291, 825, and 1,077 nuclei at 3, 4, 5.3, and 6 hpf, respectively, from 4 embryos each stage. (**B**) Quantification of Haralick Contrast as a proxy of DNA compaction in diffraction-limited images shows significant increase from 3-6 hpf. P-values determined by permutation test. (**C**) Quantification of nuclear H3K27me3 levels from Fig. 2A shows significant increase from 4-6 hpf. P-values determined by permutation test. (**D**) Expansion factor measured by nuclear cross-sectional area (linear expansion factor = 4.55; N = 894 nuclei from 6 embryos and 2,985 nuclei from 5 embryos, for unexpanded and expanded conditions). (**E**) qPCR showing the fold-change reduction in EED transcript levels following Cas13d targeting. Expression is normalized to mCherry-targeted control embryos. N = 3. P-values determined by paired t-test. (**F**) qPCR showing the fold-change reduction in EZH2 transcript levels following Cas13d targeting. Expression is normalized to mCherry-targeted control embryos. N = 3. P-values determined by paired t-test (**G**) Images of DNA corresponding to Fig. 2D. Images are maximum intensity projections of individual nuclei. (**H**) Images shown in Fig. 2F with display setting matched across all images for visual comparison. (**I**) Radial autocorrelation curves corresponding to the images in (G). (**J**) H3K9me3 staining in Cas13d EZH2- or EED-knockdown embryos compared to a non-targeting control (mCherry) at 6 hpf. N = 406, 657, and 687 nuclei from 5 embryos for EZH2, EED, and mEmerald targeted embryos, respectively. (**K**) Quantification of H3K9me3 levels from the images in (I). P-values determined by permutation test.

**Figure S3.**
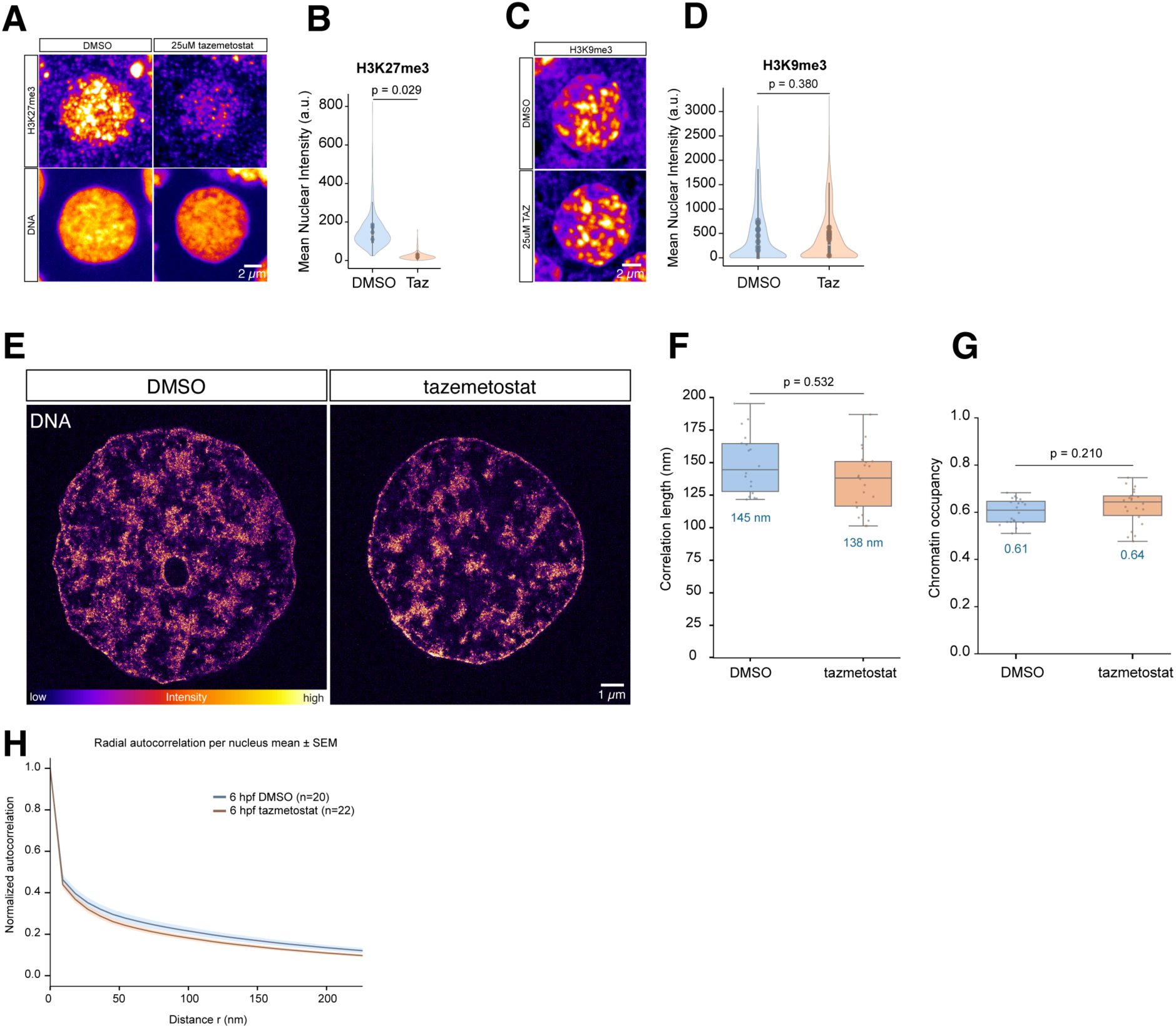
EZH2 inhibition reduces H3K27me3 without changing chromatin organization. (**A**) H3K27me3 staining in 6 hpf embryos treated with DMSO (control) or tazemetostat (EZH2 inhibitor). N = 1,103 and 791 nuclei from 4 embryos each for and DMSO and tazemetostat treated embryos, respectively. (**B**) Quantification of H3K27me3 levels from the images in (A). P-value determined by permutation test. (**C**) H3K9me3 staining in embryos treated DMSO (control) or tazemetostat at 6 hpf. N = 4,493 and 4,476 nuclei from 10 embryos each for DMSO and tazemetostat treated embryos, respectively. (**D**) Quantification of H3K9me3 levels from the images in (C). P-value determined by permutation test. (**E**) ChromExM images of individual nuclei from DMSO (control) tazemetostat (EZH2 inhibitor) treated embryos at 6 hpf. Images are shown as a single Z-slice. N = 20 and 22 nuclei from 2 and 3 embryos for DMSO and tazemetostat treatment, respectively. (**F**) Quantification of the autocorrelation length from the images in (E) shows no significant change upon EZH2 inhibition. P-value determined by permutation test. (**G**) Quantification of chromatin occupancy from the images in (E) shows no significant change upon EZH2 inhibition. P-values determined by permutation test. (**H**) Radial autocorrelation curves corresponding to the images in (E).

**Figure S4.**
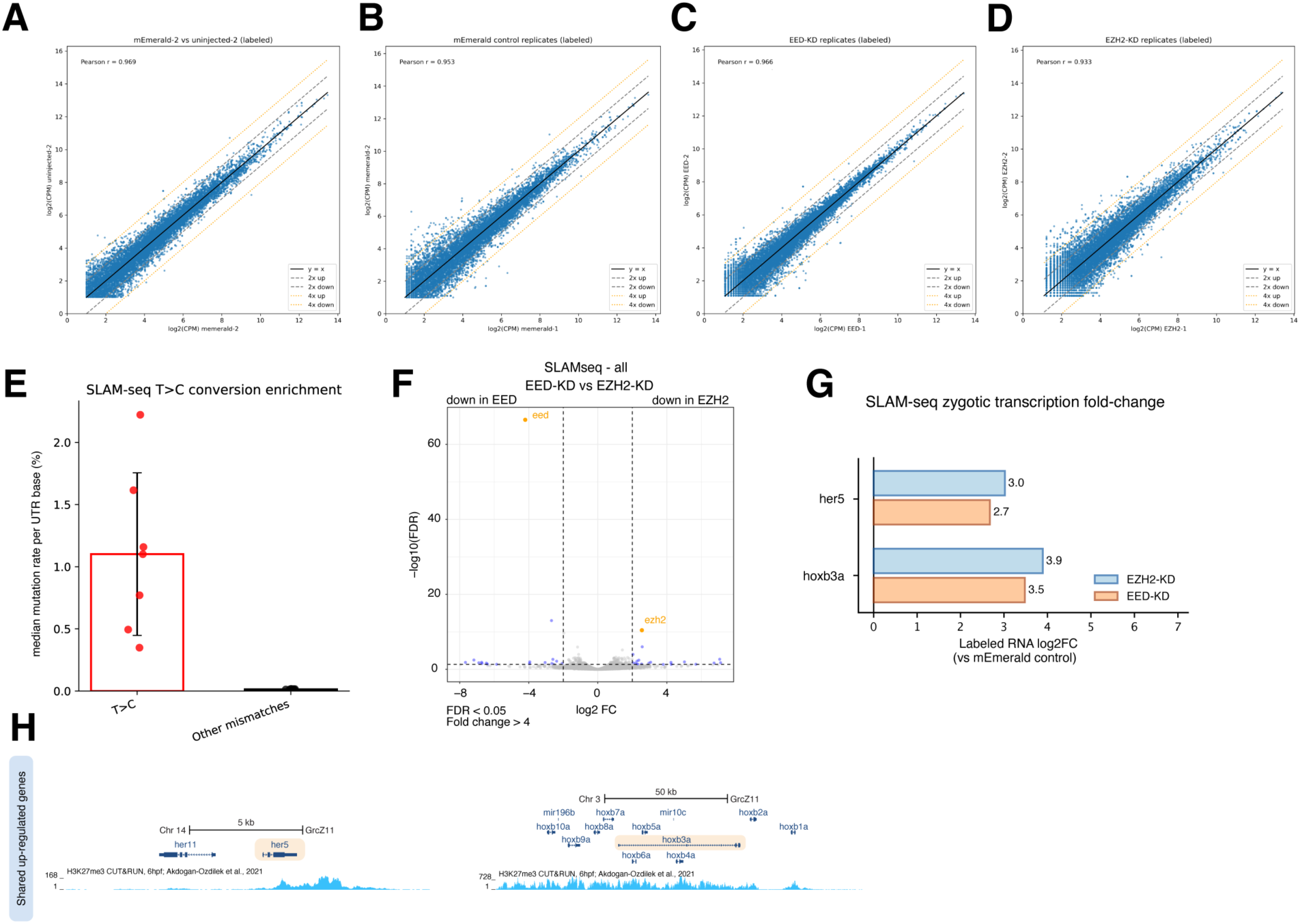
SLAM-seq shows PRC2-knockdown specificity. (**A**) Scatter plot showing the correlation for zygotic (labeled) SLAM-seq in embryos injected with Cas13d targeting mEmerald (control) and uninjected embryos. Samples are highly correlated with with Pearson R = 0.969. (**B-D**) Scatter plots showing the correlation between replicates for zygotic (labeled) SLAM-seq in embryos injected with Cas13d targeting mEmerald (control), EED, or EZH2, respectively. Replicates are highly correlated with Pearson R = 0.953, 0.966, and 0.933, respectively. (**E**) Box plot showing T>C conversion rate for all SLAM-seq samples. Each point is the median conversion rate per UTR base for one sample, with an overall median T>C conversion rate of 1.10 %. The median mismatch rate of all other possible mismatches is 0.01%. (**F**) Volcano plot showing differentially expressed genes based on maternal and zygotic (labeled and unlabeled) transcripts in EZH2 versus EED knockdown embryos. (**G**) Fold-change for select differentially expressed zygotic genes (shown in (H) in EZH2- and EED-knockdown embryos relative to mEmerald-targeted control embryos. (**H**) Genome browser tracks for genes shown in (G) showing H3K27me3 CUT&RUN (6 hpf; raw data from (*43*)) levels on significantly upregulated genes following PRC2 knockdown.

**Figure S5.**
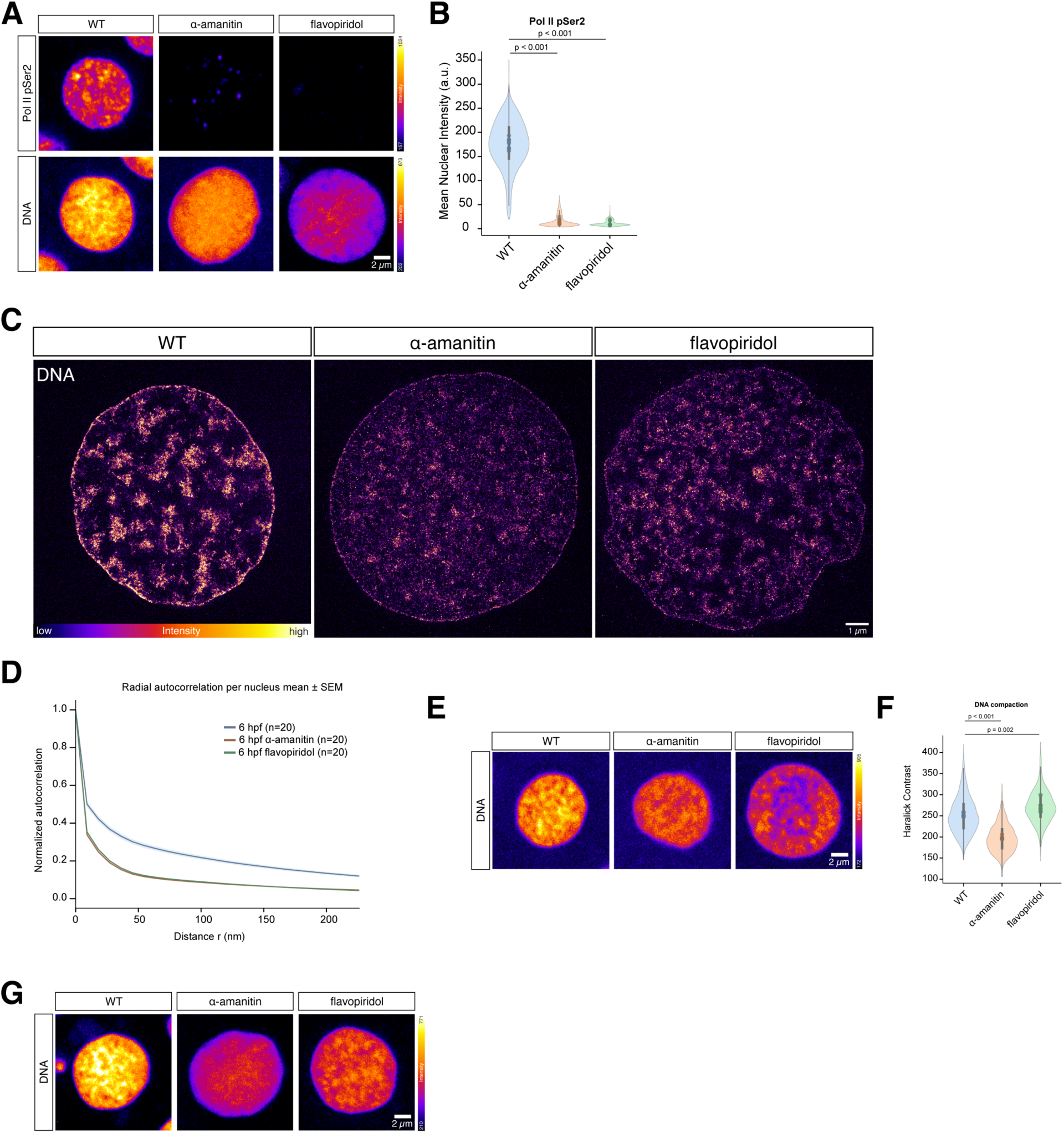
Effects of transcription inhibition on chromatin and heterochromatin marks. (**A**) Diffraction-limited immunostaining of Pol II pSer2 (elongating Pol II) shows a significant decrease following transcription inhibition. Images are maximum intensity projections of individual nuclei. N = 2,628, 1,618, and 1,189 nuclei from 8 embryos each for WT, α-amanitin, and flavopiridol, respectively. (**B**) Quantification of nuclear Pol II pSer2 levels from (A) shows a significant decrease (91% and 94% decrease for α-amanitin, and flavopiridol, respectively) upon transcription inhibition. P-values determined by permutation test. (**C**) Images shown in Fig. 3A with display setting matched across all images for visual comparison. (**D**) Radial autocorrelation curves corresponding to the images in (C). (**E**) Images of DNA corresponding to Fig. 3E. Images are maximum intensity projections of individual nuclei. N = 2,091, 1,938, and 911 nuclei from 6-7 embryos for WT, α-amanitin, and flavopiridol, respectively. (**F**) Quantification of Haralick Contrast as a proxy of DNA compaction from the diffraction-limited images in (E). P-values determined by permutation test. (**G**) Images of DNA corresponding to Fig. 3G. Images are maximum intensity projections of individual nuclei.

**Figure S6.**
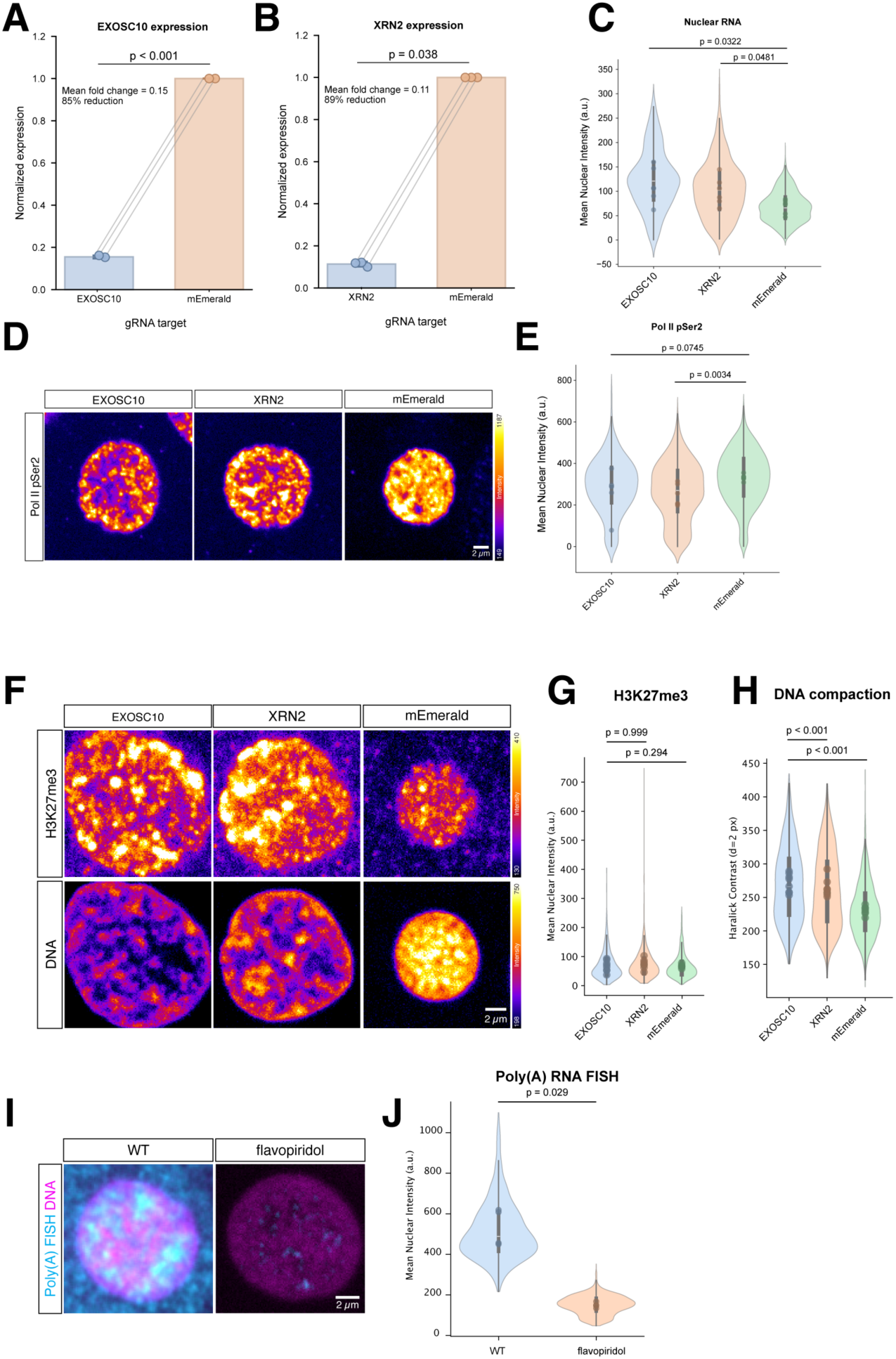
Additional characterization of RNA degradation perturbations and PolyA RNA localization. (**A**) qPCR showing the fold-change reduction in EXOSC10 transcript levels following Cas13d targeting. Expression is normalized to mEmerald targeted control embryos. N = 3 P-values determined by paired t-test. (**B**) qPCR showing the fold-change reduction in XRN2 transcript levels following Cas13d targeting. Expression is normalized to mEmerald targeted control embryos. N = 3. P-values determined by paired t-test. (**C**) Quantification of nuclear RNA levels from EU staining showin in Fig. 4A. N = 470, 526, and 1,850 nuclei from 6 embryos each for EXOSC10, XRN2, and mEmerald, respectively. P-values determined by permutation test. (**D**) Pol II pSer2 (elongating Pol II) immunostaining following EXOSC10 or XRN2 knockdown when compared to a non-targeting control (mEmerald). N = 470, 526, and 1,850 nuclei from 6 embryos each for EXOSC10, XRN2, and mEmerald, respectively. Images are max intensity projections of individual nuclei from the same embryos shown in Fig. 4A. (**E**) Quantification of the images in (D) show no significant increase in Pol II pSer2 levels following EXOSC10 or XRN2 knockdown. XRN2 knockdown leads to a mild but significant decrease in Pol II pSer2 (27% reduction). P-values determined by permutation test. (**F**) H3K27me3 immunostaining following EXOSC10 or XRN2 knockdown when compared to a non-targeting control (mEmerald). N = 1,032, 957, and 2,160 nuclei from 7-8 embryos each for EXOSC10, XRN2, and mEmerald, respectively. Images are max intensity projections of individual nuclei. (**G**) Quantification of the images in (F) show no significant change in H3K27me3 levels following EXOSC10 or XRN2 knockdown. P-values determined by permutation test. (**H**) Quantification of Haralick Contrast as a proxy of DNA compaction from the diffraction-limited images in (F) shows a significant increase following EXOSC10 or XRN2 knockdown. P-values determined by permutation test. (**I**) PolyA RNA FISH in flavopiridol treated embryos at 6 hpf. (**J**) Quantification of the images in (I) shows that transcription inhibition significantly reduces PolyA RNA levels (72% reduction). N = 1,307 and 932 nuclei from 4 embryos for WT and flavopiridol, respectively. P-values determined by permutation test.

**Movie S1. Movie through a ChromExM Z-stack from 3 hpf.**

Representative movie moving through a Z-stack of a nucleus at 3 hpf from a wild-type embryo.

**Movie S2. Movie through a ChromExM Z-stack from 6 hpf.**

Representative movie moving through a Z-stack of a nucleus at 6 hpf from a wild-type embryo.

**Movie S3. Movie through a ChromExM Z-stack from 6 hpf with α-amanitin treatment.**

Representative movie moving through a Z-stack of a nucleus at 6 hpf from an α-amanitin treated embryo.

**Movie S4. Movie through a ChromExM Z-stack from 6 hpf with flavopiridol treatment.**

Representative movie moving through a Z-stack of a nucleus at 6 hpf from a flavopiridol treated embryo

**Data S1. Cas13 sgRNA sequences.** gRNA sequences used for Cas13 targeting

**Data S2. qPCR primer sequences.** qPCR primer sequences used for measuring Cas13 target degradation

**Data S3. SLAM-seq sample information.** Information related to SLAM-seq samples

